# Tunable Low-Rate Genomic Recombination with Cre-*lox* in *Escherichia coli*: A Versatile Tool for Anoxic Environmental Biosensing and Synthetic Biology

**DOI:** 10.1101/2024.10.02.616356

**Authors:** Elisa Garabello, Hyun Yoon, Matthew C. Reid, Andrea Giometto

## Abstract

The ability to induce heritable genomic changes in response to environmental cues is valuable for environmental biosensing, for experimentally probing microbial ecology and evolution, and for synthetic biology applications. Site-specific recombinases provide a route to genetic memory via targeted DNA modifications, but their high specificity and efficiency are offset by leaky expression and limited tunability in prokaryotes. We developed a tightly regulated, titratable Cre recombinase system for *Escherichia coli* that achieves low recombination rates and minimal basal activity. Implemented on both plasmids and the chromosome, the latter showed superior retention of genetic memory across generations. These features make the system broadly useful for environmental biosensing and other applications. To demonstrate applicability to environmental biosensing, we developed a whole-cell recombination-based biosensor for arsenite, a toxic and ubiquitous pollutant that is primarily mobilized in anoxic environments such as flooded soils, sediments, and aquifers. However, existing arsenite whole-cell biosensors face limitations in sensitivity and workflow in anaerobic settings. Our biosensor reliably recorded anoxic arsenite exposure as a stable genetic memory for delayed fluorescence readout in aerobic conditions, with detection sensitivity comparable to conventional wet chemical methods. By decoupling exposure from measurement, this approach offers a foundation for arsenite biosensing under field-relevant conditions, including redox variability and other physicochemical gradients, without the constraints of anoxic measurement. More broadly, the ability to induce low-rate, heritable genetic changes expands the genetic toolkit for environmentally responsive systems, with applications in environmental monitoring, bioproduction, bioengineering, as well as experimental studies of microbial ecology, evolution, and host-microbe interactions.

**IMPORTANCE:** Arsenic is a toxic and globally prevalent pollutant, mobilized primarily under anoxic conditions where detection is challenging. Whole-cell biosensors offer a promising route for monitoring bioavailable arsenic *in situ*, but their development has largely focused on aerobic conditions, with anoxic assays limited by sensitivity and workflow constraints. Genetic tools that enable heritable, low-frequency genomic changes in bacteria can expand biosensor capabilities by recording transient exposures and supporting applications in environmental monitoring, synthetic biology, and quantitative microbial population dynamics research. Here, we developed a tightly regulated, chemically inducible Cre-*lox* system in *Escherichia coli* that enables recombination at low, tunable rates. We demonstrate its utility by constructing an arsenite biosensor that reliably detects low concentrations and records exposures under both aerobic and anoxic conditions. This approach is broadly applicable for biosensors designed for field deployment and for experiments investigating microbial ecology and evolution, where controllable genetic diversification may be desirable.

## INTRODUCTION

Controlling microbial responses to environmental signals is central to developing biosensing strategies suited to complex environmental settings (1) and to developing and validating quantitative models of microbial ecology and evolution (2–4). Environmental biosensors designed to detect chemical contaminants would benefit from systems capable of stably recording transient exposures, particularly in field conditions where continuous monitoring is impractical. Similarly, tools that enable phenotypic or genotypic diversification at controlled rates can be used to simulate mutational processes (3), track lineages, engineer controlled diversification (2), or probe community responses to controlled perturbations. All of these can be used to test ecological and evolutionary theory. Both use cases require genetic systems that can induce heritable changes at low, tunable rates in response to specific environmental inputs.

Recombinase-based systems offer a powerful approach for designing environmental biosensors (5,6) because site-specific recombination produces permanent, heritable DNA changes that can serve as a genetic record of transient environmental exposure without the need for continuous signal input. This capability is particularly valuable for field deployment, where exposures may occur under conditions that are challenging for real-time measurement (e.g., anoxic environments or fluctuating physicochemical gradients) and where sample analysis often takes place well after collection. For example, a recombinase-based biosensor for arsenite could detect exposure events in anoxic settings relevant to arsenic mobility (7) and allow the record to be read out later under aerobic conditions, avoiding the need for specialized anaerobic measurement setups (8). These practical advantages, combined with the ability to tune recombination rates, make recombinases well-suited for environmental monitoring and other applications. To realize this potential, it is important to understand and overcome their current limitations in bacterial systems.

Site-specific recombinases and nucleases enable heritable genetic changes by allowing targeted genome modifications at specific loci *in vivo*. These enzymes can be used to induce reversible or irreversible phenotypic changes, often coupled to reporter systems such as fluorescent proteins. Among them, Cre is a tyrosine recombinase that recognizes two short DNA sequences, called *lox* sites, and catalyzes recombination between them with high specificity and efficiency. When two *lox* sites are oriented in the same direction in the same DNA molecule, Cre excises the DNA between them (9,10). Depending on the configuration of *lox* sites, Cre can also mediate DNA inversion, integration, or translocation, expanding its utility in genetic circuit design. The resulting genotype is heritable and maintained even after Cre expression is interrupted. This feature, shared by other recombinases such as Flp and Bxb1, has been exploited in synthetic genetic logic gates and circuits with a memory component (6,11). Unlike nucleases such as CRISPR-Cas, recombinases do not rely on host double-strand break repair machinery, allowing them to carry out precise recombination events autonomously. This characteristic is particularly advantageous in prokaryotes like *E. coli*, where DNA repair options, though varied, predominantly depend on homology-directed mechanisms that may be less efficient or more error-prone. Because recombinase-mediated DNA changes are permanent and do not require ongoing expression, Cre*-lox* systems offer a robust means of encoding long-term memory of environmental inputs.

Despite these advantages, applying recombinase systems like Cre in *E. coli* remains challenging, particularly in scenarios that require precise control over recombination rates. In bacteria, recombinases have direct access to chromosomal and plasmid DNA, and even minimal basal expression can lead to rapid and irreversible switching in a large fraction of the population. This makes it difficult to implement systems that require slow, partial, or tunable transitions, which are valuable for experiments involving controlled genotypic diversification (2), lineage tracing (3), or temporally integrated biosensing (6,12,13). In eukaryotic hosts, recombinase activity can often be tightly regulated through compartmentalization or ligand-dependent nuclear import. For instance, the yeast Mother Enrichment Program uses β-estradiol to control Cre nuclear import via a fusion with a hormone-binding domain, enabling tight regulation of recombinase activity (14). In contrast, *E. coli* lacks these regulatory layers, making transcriptional leakiness a critical issue. Recent work by Williams *et al.* (15) demonstrated that the effective rate of Bxb1-mediated differentiation can be tuned by controlling the integrase expression levels. Most existing implementations of Cre have been optimized for rapid and complete switching, leaving a need for systems that support persistent, low-frequency recombination under precise regulatory control. To control Cre recombinase activity in bacteria, Sheets *et al.* (16,17) constructed and characterized different variants of an optogenetic system that regulates it at the post-translational level. Although using a light signal to regulate Cre activity brings benefits such as temporal control over recombination, it also limits the experimental setup that can be employed, due to the necessity of controlling sample illumination at all times. These illumination requirements restrict deployment outside the lab and make the approach ill-suited to applications such as environmental biosensing.

Here, we developed and characterized titratable Cre-*lox* reporters with low background expression in *E. coli*. By regulating Cre at both the transcriptional and post-translational levels and implementing these systems either on plasmids or on the chromosome, we enable phenotypic switching at low, chemically inducible rates. We deliberately constrained the system so that, over a typical experiment (∼10 generations), only a fraction of cells converts to the recombined genotype when exposed to analyte concentrations within the range of interest. Under low-rate conditions, small differences in inducer concentration produce small differences in recombination rates, which accumulate over time and are amplified into detectable differences in the fraction of recombined (fluorescent) cells. This enables sensitive detection and expands the range of inducer concentrations that can be resolved. As a result, these systems are suitable for a range of uses beyond traditional synthetic biology applications, such as metabolic engineering and long-duration biosensing, and extends to experiments in cell physiology, ecology, and evolutionary biology, where controlled genotypic diversification needs to be maintained over many generations.

We demonstrate the broad applicability and advantages of these constructs by developing a Cre-recombinase-based biosensor for quantification of bioavailable arsenite and compare its performance to more conventional transcription-based fluorescent reporters. Arsenic (As) is a widespread environmental contaminant that poses major risks to both human health and ecological systems. The U.S. Environmental Protection Agency (EPA) adopted the World Health Organization (WHO) drinking water Maximum Contaminant Level (MCL) for arsenic maximum contaminant limit (MCL) of 10 μg/L. Despite this regulation, it is estimated thant more than 6 million people in the U.S. are exposed to higher concentrations exceeding this threshold (7). In regions such as Bangladesh, where groundwater arsenic contamination is widespread, the limit set at 50 μg/L (18). Arsenic sensors for aqueous samples must therefore reliably detect concentrations spanning the regulatory limit as well as the higher levels commonly observed in contaminated groundwater. Although average freshwater arsenic concentrations are approximately On average arsenic contamination in freshwater at 0.8 μg/L, reported values span more than four orders of magnitude, reaching 600 μg/L and, in extreme cases, up to or even 13 mg/L (19). In anaerobic environments such as waterlogged rice paddies, sediments, and groundwater aquifers, microbial processes mobilize arsenic primarily in its trivalent form, arsenite (As(III)), which is highly toxic and mobile. Detecting arsenite under these conditions is critical for understanding its biogeochemical cycling (7) and assessing exposure risks. While many arsenic whole-cell biosensors have been developed (Table S1), relatively few can operate under anoxic conditions (20). These biosensors exhibit lower sensitivity than systems employing oxygen-dependent fluorescent reporters (21,22), largely due to the reduced brightness of flavin-based fluorescent proteins. Moreover, they require fluorescence measurements to be performed within anaerobic chambers, or potentially *in situ* under anoxic conditions. These constraints make their deployment more challenging than measurements conducted in oxic conditions after exposure. To address this gap, we engineered a Cre-recombinase-based biosensor that can record transient arsenite exposure under anoxic conditions, preserve this information in the genome, and allow delayed readout under oxic conditions.

## RESULTS

This section is organized as follows: First, we describe the optimization of Cre-*lox* constructs for titratable, low-rate recombination by expressing *cre* from a promoter inducible by 4-isopropylbenzoic acid (cuminic acid), supplied as its cumate salt. Second, we characterize the inducer response of these constructs at both the population and single-cell levels. Third, we present the development of a Cre-recombinase-based arsenite biosensor and assay its behavior in aerobic and anoxic growth conditions. Finally, we examine the broader applicability of our constructs and test the reporters for potential loss of genetic memory or fluorescence readout over multiple generations.

Aiming to reduce and control Cre activity, our first construct mounted on a low copy number plasmid (p*-cymR-cre*[LVA]-loxPP*-eyfp)*, with origin of replication p15A as all other plasmid unless otherwise noted) expressed *cre* from the cumate-sensitive promoter P*_cymRC_* obtained from the “Marionette” sensors collection (23,24). This promoter is repressed by the regulator CymR^AM^ (constitutively expressed from the chromosome of the host strain Marionette-Clo, sAJM.1504 (23)) in the absence of cumate. Both *cymR^AM^* and P*_cymRC_* are optimized to provide lower leakage and increased dynamic range (23). In addition, this construct includes *cymR^AM^*, which maintains the ratio between the copy numbers of the regulator *cymR^AM^* and the promoter P*_cymRC_* close to one. Finally, we added the C-terminal ssrA-LVA degradation tag (25) to *cre* to reduce Cre half-life in the cell and thus its activity (Fig. 1).

**Fig. 1.**
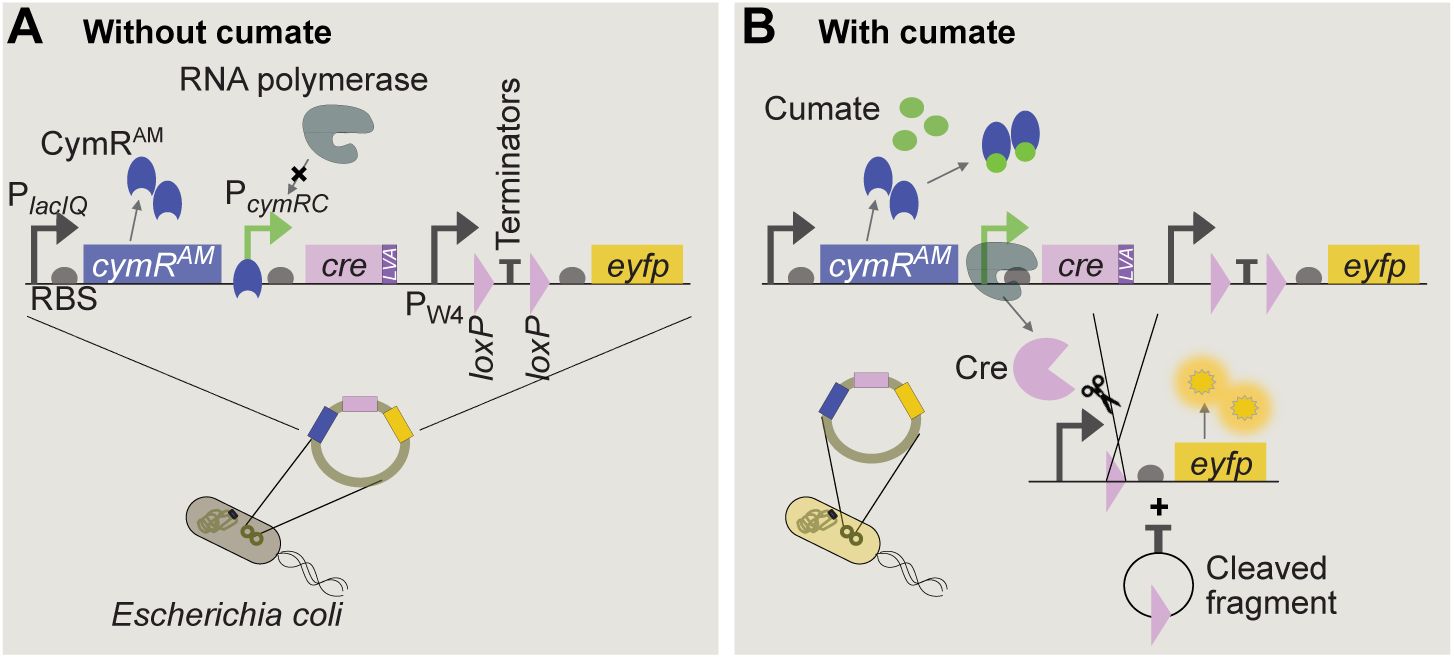
Schematic of the Cre-*loxP* reporter system in *E. coli* on a plasmid. (A) In the absence of cumate, the constitutively expressed repressor CymR^AM^ occupies the CuO operator sites in P*_cymRC,_* inhibiting *cre* transcription. (B) When cumate is present, it binds to the repressor, which detaches from the operator site allowing *cre* transcription. If the *loxP* sites are positioned in the same orientation, Cre excises the terminators located between the *loxP* sites and transcription of *eyfp* is initiated.

To test the inducer-response curve of this circuit and other variants, we assembled a construct, loxPP, composed of the constitutive promoter P_W4_ (16) and two transcriptional terminators (TT, see Table 1) flanked by two *loxP* sites (part of phage P1’s wild-type Cre-*loxP* system) with the same orientation, followed by the enhanced yellow fluorescent protein gene (*eyfp*). In the presence of both *loxP* sites, the two transcriptional terminators interrupt transcription, leaving *eyfp* unexpressed (Fig. 1A). Upon Cre-induced recombination of the two *loxP* sites, the transcriptional terminators are excised (Fig. 1B), enabling expression of the fluorescent protein that we detected using a fluorescence plate reader or a flow cytometer, depending on the assay. Simultaneous measurements of optical density at 600 nm (OD600) and fluorescence at the plate reader allowed us to identify cumate concentrations that did not significantly affect growth rates (Table S2), while promoting strong Cre activity.

**Table 1.** Summary of relevant plasmids and strains used in this study.

| Strain | Source | Genotype |
| --- | --- | --- |
| sAJM.1504 | Meyer *et al.* (23) | *E. coli* Marionette-Clo (DH10B background), *cymR<sup>AM</sup>* on chromosome |
| bAG44 | Pothier *et al.* (30) | *E. coli* NEB10-beta, (DH10B background), host of As biosensor |
| c-*cre*[LVA]-loxPP-*syfp2* (bEG57, chromosomal reporter) | This work | sAJM.1504; *cymR<sup>AM</sup>* // *attTn7::P<sub>cymRC</sub> – cre*[LVA] – T – *P<sub>rpsL</sub> – loxP – TT – loxP – syfp2* |
| Plasmid | Source | Key Inserts (p15A origin unless otherwise noted) |
| pAJM.657 | Meyer *et al.* (23) | *cymR<sup>AM</sup>*, *P<sub>cymRC</sub> – eyfp* |
| pAJM.717 | Meyer *et al.* (23) | *kanR*, *P<sub>Tet</sub> – eyfp* |
| p-*cymR-cre*[LVA]-lox511P-*eyfp* (pAG56) | This work | *cymR<sup>AM</sup>*, *P<sub>cymRC</sub> – cre*[LVA] – T – *P<sub>W4</sub> – lox511 – TT – loxP – eyfp* |
| p-*cymR-cre*[LVA]-loxPP-*eyfp* (pEG36, plasmid reporter) | This work | *cymR<sup>AM</sup>*, *P<sub>cymRC</sub> – cre*[LVA] – T – *P<sub>W4</sub> – loxP – TT – loxP – eyfp* |
| p-*cymR-cre*-loxPP-*eyfp* (pEG37) | This work | *cymR<sup>AM</sup>*, *P<sub>cymRC</sub> – cre* – T – *P<sub>W4</sub> – lox511 – TT – loxP – eyfp* |
| p-*cre*[LVA]-loxPP-*eyfp* (pEG39) | This work | *P<sub>cymRC</sub> – cre*[LVA] – T – *P<sub>W4</sub> – loxP – TT – loxP – eyfp* |
| p-*cre*[LVA]-loxPP-*syfp2* (pEG55) | This work | *P<sub>cymRC</sub> – cre*[LVA] – T – *P<sub>rpsL</sub> – loxP – TT – loxP – syfp2* |
| pMP01 | Pothier *et al.* (30) | ArsRBS2 – *mCherry* (pUCP19 backbone) |
| p-ArsRBS2-<br>*cre*[LVA]-<br>*loxPP-syfp2*<br>(pEG69) | This work | ArsRBS2 – *cre*[LVA] – T – $P_{rpsL}$ – *loxP* – TT – *loxP* – *syfp2* |
| p-ArsRBS2-<br>*syfp2*<br>(pEG78) | This work | ArsRBS2 – *syfp2* |
| As negative<br>control<br>(pEG125) | This work | T – $P_{rpsL}$ – *loxP* – TT – *loxP* – *syfp2* |
| **Units** | **Source** | **Genetic Elements** |
| T | Synthesized by<br>Twist<br>Bioscience,<br>sequence from<br>Chen *et al.* (43) | Transcriptional terminators L351P13-BBa_B0062-R-L3S3P22 |
| TT | Synthesized by<br>Twist<br>Bioscience,<br>sequence from<br>Chen *et al.* (43) | Transcriptional terminators *ECK120033736* – *L3S2P11* |
| ArsRBS2 | Pothier *et al.* (30) | [ArsR binding site] – $P_{arsR}$ – *arsR* – [ArsR binding site] |
| *loxPP* | This work | *loxP* – TT – *loxP* |
| *lox511P* | This work | *lox511* – TT – *loxP* |

Fig. 2A shows that this system (p-*cymR-cre*[LVA]-loxPP*-eyfp*) has low leakage and titratable Cre activity across a reproducible range. To further vary the titrability range of the construct, we tested different regulatory strategies at both the transcriptional and the post-translational level (Fig. 2B-C). Specifically, we explored if we could alter the activity of our low-rate Cre-*loxP* system by reducing the gene copy number of *cymR^AM^*, by removing it from the plasmid (p*-cre*[LVA]-loxPP*-eyfp*), by adopting heterologous *lox* sites (p*-cymR-cre*[LVA]-lox511P*-eyfp*), or by removing the Cre degradation tag (see additional notes in Supplemental Materials). Recombination rates increased, as expected, when the regulator gene *cymR^AM^* was removed from the plasmid (p*-cre*[LVA]-loxPP*-eyfp*, Fig. 2B), leaving only the chromosomal copy. This may be of interest for applications that require a reporter with high sensitivity and quick response times. Conversely, we observed a dramatic drop in the fraction of fluorescent cells in the population, and thus in Cre activity, when the first wild-type *loxP* was replaced with *lox511* (p*-cymR-cre*[LVA]-lox511P*-eyfp*, Fig. 2C), which differs from *loxP* by a single nucleotide in the spacer, the central region of *lox* sites that determines recombination specificity and directionality (26,27). We were able to obtain higher fractions of fluorescent cells by inducing the construct with 50 and 100 μM cumate, but observed a sizable fitness cost (Table S2). Since this construct had a reduced titrability range compared to other variants, we did not characterize it further. Still, it may be of interest for longer assays, for example for growth in chemostats in which recombination of homologous *lox* sites constructs such as p*-cymR-cre*[LVA]-loxPP*-eyfp* would likely reach saturation (100% fluorescent cells) in a short time. Removal of Cre degradation tag resulted in frequent recombination even in the absence of the inducer, making it impossible to obtain clonal populations with two intact *loxP* sites, unless heterologous *lox* sites were used. The various plasmid variants are listed in Table 1 (Materials and Methods).

**Fig. 2.**
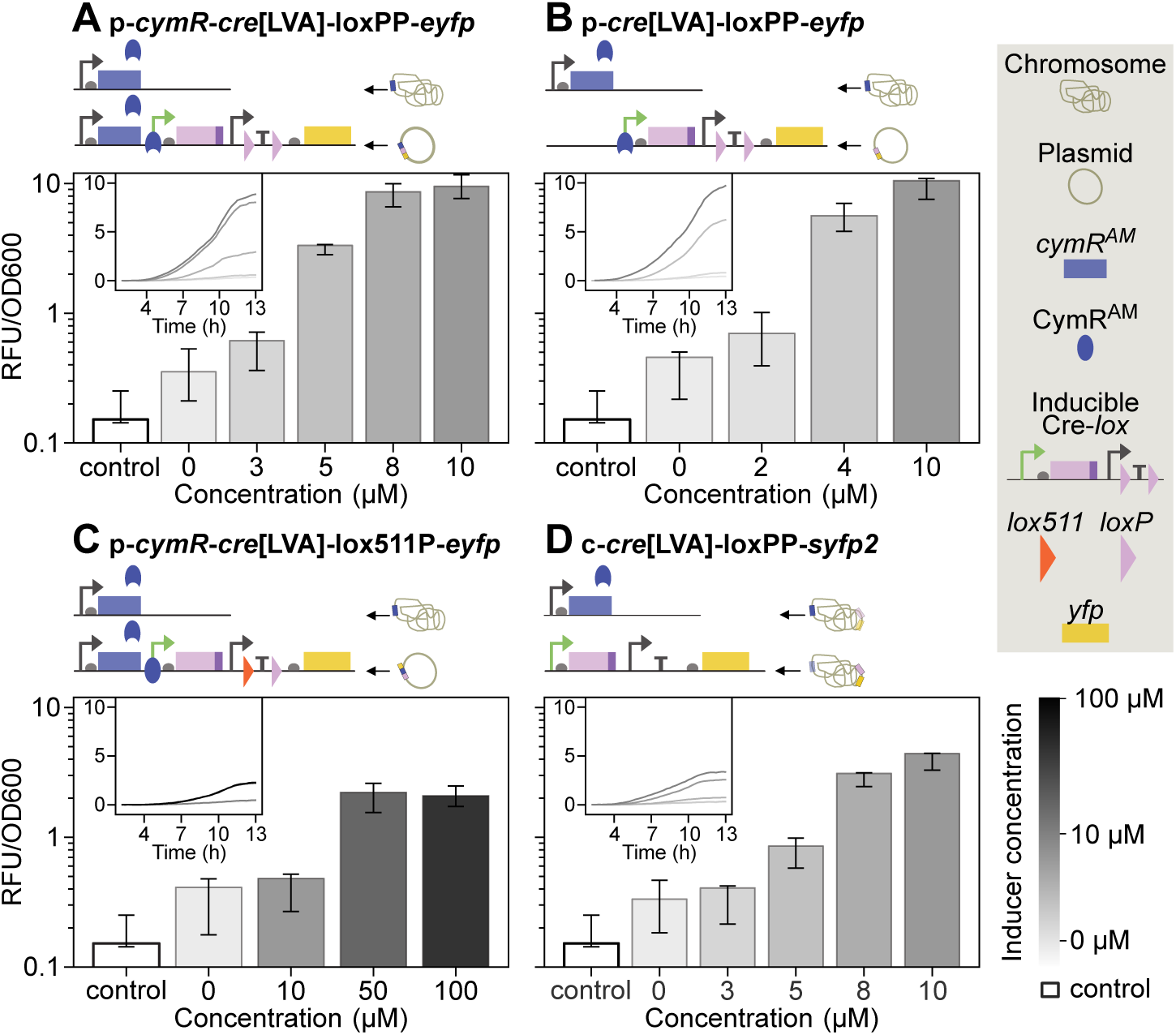
Inducer-response curves showing the relative fluorescence output of four Cre-*lox* reporter variants measured at theusing a fluorescence plate reader. (A-D) Relative fluorescence divided by optical density at 600 nm (OD_600_) (see Materials and Methods) after 10 hours from inoculation. The range of titrability varied with the copy number of the repressor gene *cymR^AM^* (A vs B), the identity of *lox* sites (homologous in A vs heterologous in C), and the genomic location of the circuit (plasmid in A-C vs chromosome in D). Relative fluorescence values varied with the same factors and with reporter design, as plasmid and chromosomal reporters use different fluorescent proteins and associated promoters and RBS. The negative control is the fluorescence intensity recorded for the parent strain Marionette-Clo which does not contain any Cre-*lox* circuit, nor any fluorescent protein. All strains have one copy of *cymR^AM^* on the chromosome. Technical replicates were averaged within each biological replicate to produce a mean estimate per experiment. Insets show the median of these means as relative fluorescence output vs time, without OD_600_ normalization. Bars and insets show the median of these means, with error bars indicating the full range (min-max) across biological-replicate estimates. For p*-cre*[LVA]-loxPP*-eyfp*In panel B, only two biological replicates were included for intermediate concentrations 2 μM and 4 μM due to a setup error in the third replicate, which used incorrect inducer concentrations.

After constructing the plasmids described above and characterizing their inducer-response curves, we integrated a Cre-*loxP* reporter sequence with homologous *loxP* sites at *attTn7*, generating strain c-*cre*[LVA]-loxPP*-syfp2*. The integrated sequence was amplified from a plasmid encoding *cre*[LVA] and *syfp2* (p*-cre*[LVA]-loxPP*-syfp2*), which was derived from p*-cre*[LVA]-loxPP*-eyfp* by replacing *eyfp* with the brighter yellow super fluorescent protein gene *syfp2* and by using a stronger promoter (P*_rpsL_*) and ribosome binding site to increase the fluorescence signal. Chromosomal integration guarantees a more homogeneous gene copy number in the population and longer stability of the Cre-*loxP* reporter construct across generations, even in the absence of the selective antibiotic, as demonstrated below.

We decided to further characterize reporters of interest at the single-cell level using flow-cytometry, as we reasoned that it would allow us to detect recombination events earlier than with the fluorescence plate reader. We identified the plasmid p*-cymR-cre*[LVA]-loxPP*-eyfp* as a suitable reporter for quantitative characterization of inducer-response curves, as it exhibited a broad titrability range at relatively low inducer concentrations. At 10 μM cumate, the construct reached saturation within a single growth curve, as evidenced by all cells exhibiting fluorescence. In parallel, we characterized the chromosomally integrated reporter c-*cre*[LVA]-loxPP*-syfp2*. We selected p*-cymR-cre*[LVA]-loxPP*-eyfp* as the plasmid counterpart for comparison with the chromosomal reporter because the two constructs have approximately matched gene copy numbers of *cymR^AM^* and *cre*[LVA]. Hereafter, we focus exclusively on these two constructs and refer to p*-cymR-cre*[LVA]-loxPP*-eyfp* as the *plasmid reporter* and to c-*cre*[LVA]-loxPP*-syfp2* as the *chromosomal reporter*.

To capture the kinetics of these reporters, we collected samples of the cell population during the growth phase (Fig. 3A). Cytometry measurements allowed us to carry out a high-throughput analysis of the fraction of cells expressing the fluorescence reporter and to observe fluorescence heterogeneity at the single cell level (Fig. 3D), even when the population fluorescence was well below the detection limit of the plate reader. In addition, we performed colony PCR on samples treated with different inducer concentrations and collected at various time-points to confirm the presence of both the original (two *loxP*) and recombined (one *loxP)* sequences in populations that appeared mixed by flow-cytometry (Fig. S2). To estimate recombination rates (Fig. 3C), we fit a logistic or exponential curve (Materials and Methods) to the fraction of cells with one *loxP* measured with flow-cytometry and report the corresponding rate parameters in Table S3 (see also Fig. S1).

**Fig. 3.**
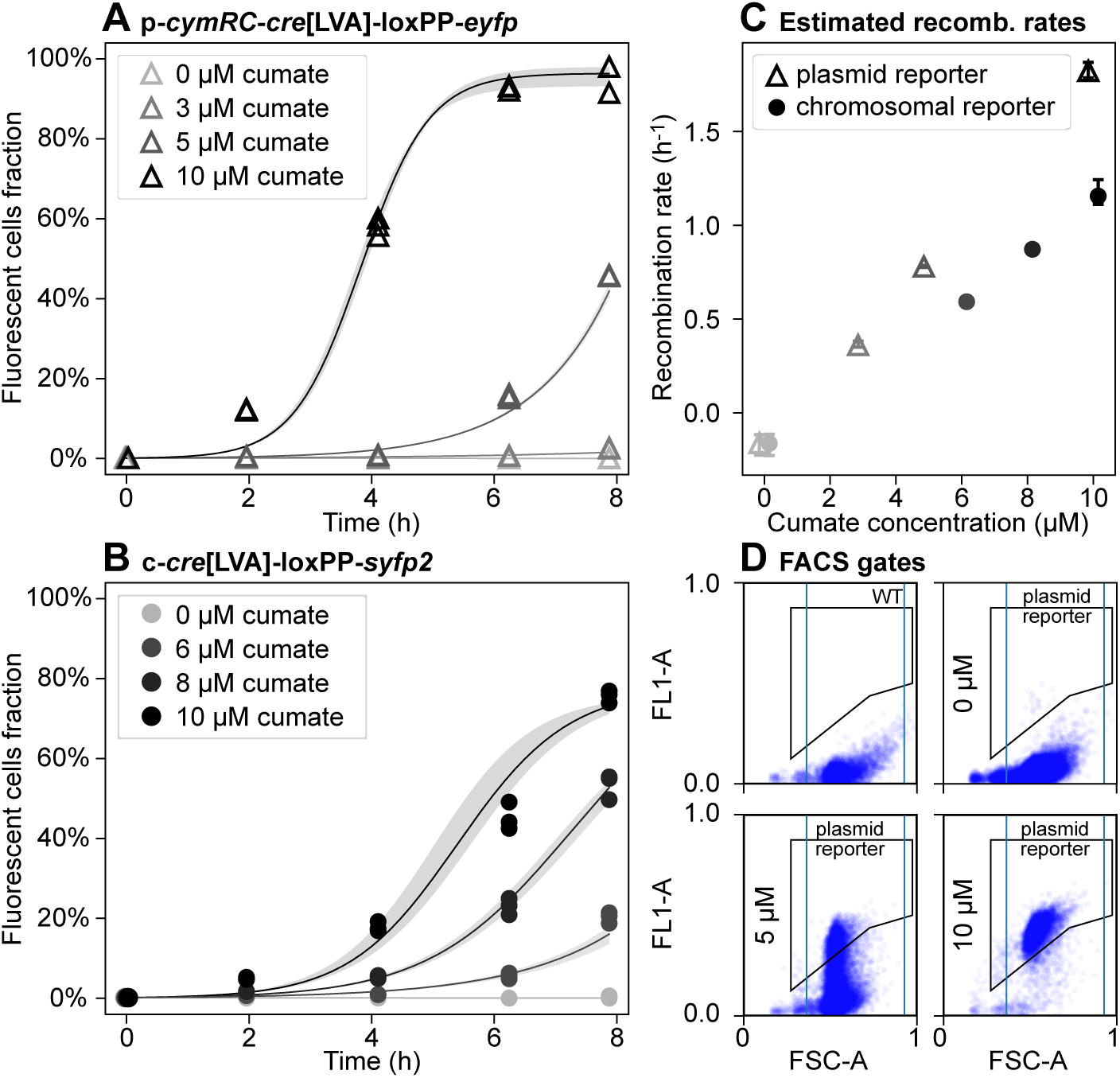
Inducer-response curves for the low-rate Cre-*loxP* plasmid p*-cymR-cre*[LVA]-loxPP*-eyfp* and chromosomal c*-cre*[LVA]-loxPP*-syfp2* reporter systems measured at the flow-cytometer. (A-B) Fraction of fluorescent cells in the population with the plasmid (A) or chromosomal (B) reporter. Each point represents a technical replicate. Time point 0 h is the same for all treatments and is the fraction of fluorescent cells in the starting culture after exiting the lag phase and before adding the inducer. Solid curves show the mean of the best-fit model predictions (Materials and Methods) across technical replicates; shaded bands indicate the min-max envelope of the replicate-specific best-fit curves. Corresponding plots for two additional biological replicates are reported in Fig. S1. (C) Best-fit estimates of recombination rates as a function of the inducer concentration. Error bars show the minimum and maximum estimated rates across technical replicates. (D) Example flow-cytometry data for plasmid reporter samples, after six hours of exposure to cumate, showing single-cell fluorescence signal area (FL1-A) versus forward-scatter area (FSC-A), in arbitrary units. Threshold gates are indicated by blue vertical lines (FSC-A channel) and black polygons (FL1-A), with lower boundaries defined based on the control strain. The gates were defined manually, with the lower boundary drawn to exclude the control strain.

To demonstrate the applicability of these designs for environmental biosensing, we built a proof-of-concept Cre-*loxP* reporter in which *cre* transcription is induced by arsenite. To this end, we expressed both *cre*[LVA] and *E. coli*’s transcriptional repressor *arsR* (28) from the promoter P*_arsR_* in a bicistronic design. We refer to this reporter as p-ArsRBS*2-cre*[LVA]-loxPP*-syfp2,* where ArsRBS2 (Fig. 4A) denotes a regulatory module comprising P*_arsR_* and the *arsR* gene flanked by two ArsR binding sites (29,30). We grew this recombinase-based arsenite-sensitive reporter in the defined medium MOPS Glycero-Phosphate (MGP), with concentrations of As(III) as low as 15 nM, comparable to the sensitivity of existing whole-cell arsenic biosensors and Gutzeit method-based field-test kits (18,31). To compare the sensitivity of our recombinase-based reporter p-ArsRBS*2-cre*[LVA]-loxPP*-syfp2* to that of a transcription-based reporter, we used a modified version of the plasmid reporter pMP01 from Pothier *et al.* (30), p-ArsRBS2*-syfp2*, carrying *syfp2* instead of *mCherry*, with the same backbone as p-ArsRBS*2-cre*[LVA]-loxPP*-syfp2* (Fig. 4B). Unlike p-ArsRBS2*-syfp2* where *syfp2* transcription is regulated via the promoter P*_arsR_*, expression of *syfp2* in p-ArsRBS*2-cre*[LVA]-loxPP*-syfp2* is regulated by a strong, constitutive promoter. Note that the translation rates of Cre and SYFP2 differ in the two constructs due to the use of different ribosome binding sites. To compare the two biosensors, we took endpoint, single-cell fluorescence measurements at the flow-cytometer for both reporters, grown with arsenite concentrations from 0 to 60 nM in MGP. Since the transcription-based reporter returns a continuous fluorescence response, we took the median single-cell fluorescence intensity as the representative measure of the overall expression level. We found that both the p-ArsRBS2-*cre*[LVA]*-*loxPP*-syfp2* and p-ArsRBS2*-syfp2* reporters could detect 15 nM arsenite (*P*-value < 0.05), which is lower than the EPA threshold values (31) for arsenic contamination in drinking water (10 ppb As, equivalent to ∼133 nM As). We observed limited leakage in p-ArsRBS*2-cre*[LVA]-loxPP*-syfp2*, with the assay performed in the absence of arsenite yielding an additional 0.11% recombined cells (median across experiments). This basal expression was expected, due to ArsR-mediated negative feedback on P*_arsR_*, which drives the bicistronic *arsR*-*cre* transcriptional unit. By comparison, in the treatments with added arsenite, the median fraction of newly recombined fluorescent cells across biological replicates ranged from 0.21% to 0.71%, corresponding to the minimum and maximum inducer concentrations, respectively.

**Fig. 4.**
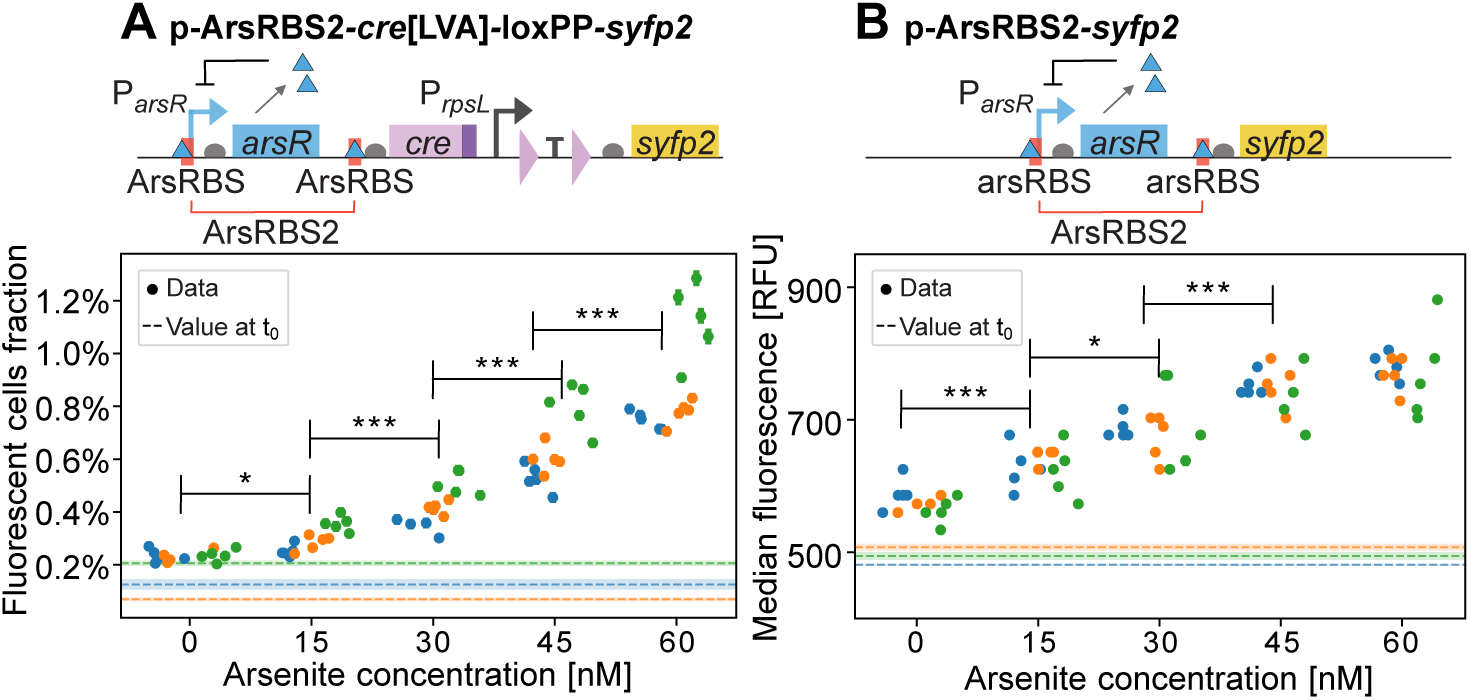
Schematics of arsenite whole-cell biosensors and endpoint fluorescence measurements collected at the flow-cytometer. (A) Recombinase-based biosensor schematics (top) and fraction of fluorescent cells at the end of the assay (bottom; error bars report the binomial standard error). (B) Transcription-based biosensor schematics (top) and median cell fluorescence in each sample (bottom); standard errors computed via bootstrapping are smaller than symbols. In both panels, each point is a technical replicate, with three biological replicates shown in different colors. The horizontal dashed lines represent the initial fluorescent cells fraction for each biological replicate, before adding the inducer, with shaded areas around them showing the associated binomial error on the fractions. The third biological replicate (green symbols) was incubated an hour longer (18 vs 17 hours) with respect to the two other replicates, which might explain the larger fraction of fluorescent cells compared to the other two experiments. Note that due to different designs, the measured response is different depending on the bioreporter: for the recombinase-based biosensor, we report on the y-axis the fraction of fluorescent cells, counted by cytometry gating; for the transcription-based reporter, we report the median cell fluorescence intensity.

The background expression of the recombinase-based arsenite bioreporter may arise from leakiness of the P*_arsR_* promoter, which would lead to low-level expression of Cre and unintended recombination, or from inefficient termination by the double transcriptional terminators flanked by the *loxP* sites. To distinguish between these two potential sources of background signal, we compared the fluorescence of cells harboring p-ArsRBS*2-cre*[LVA]-loxPP*-syfp2* grown in MGP without arsenite to that of a negative-control plasmid obtained by deleting the ArsRBS2*-cre*[LVA] regulatory module and retaining only the backbone and the construct P*_rpsL_-loxP-*TT*-loxP-syfp2* (Fig. 5 and Fig. S3). The negative control displayed a higher median single-cell fluorescence than the parental strain (Fig. S3), which lacks any fluorescent reporter gene, indicating a small amount of basal expression from the constitutive promoter P*_rpsL_*. At the same time, the biosensor exhibited fluorescence levels above the control baseline, supporting the hypothesis that leaky Cre expression causes a small number of *loxP* pairs to recombine even in absence of arsenic.

**Fig. 5.**
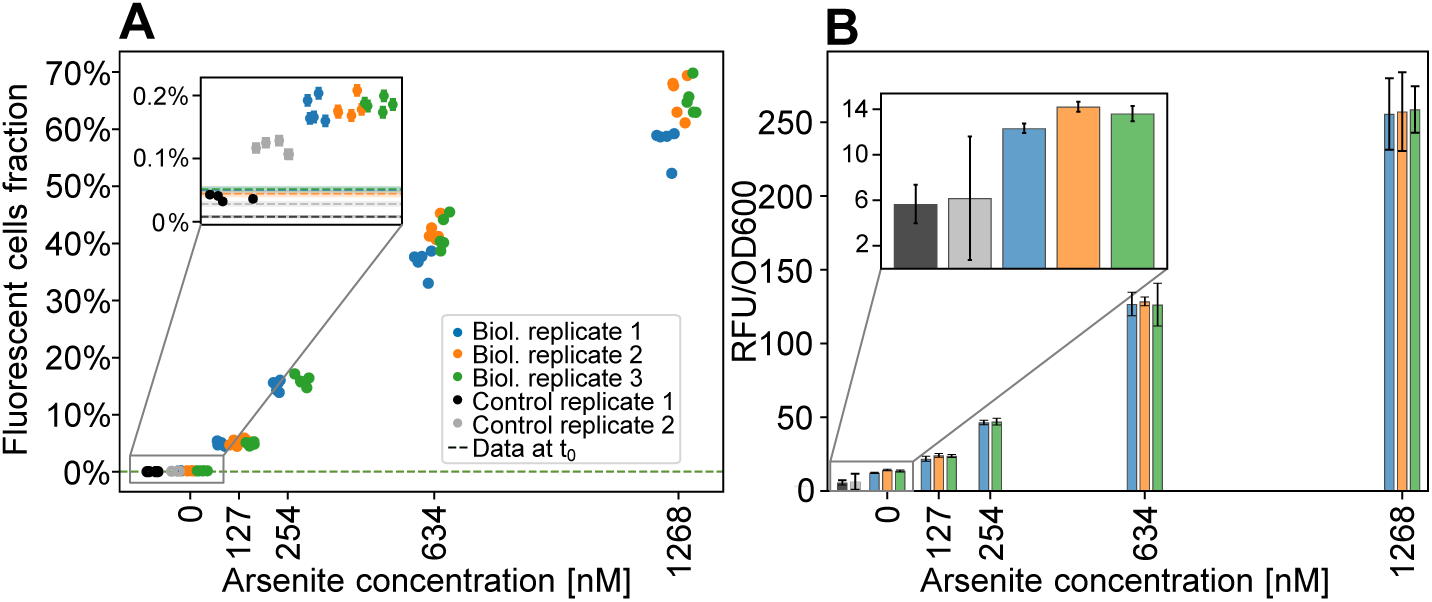
Endpoint fluorescence measurements of the recombinase-based biosensor p-ArsRBS*2-cre*[LVA]-loxPP*-syfp2* following exposure to arsenic concentrations ranging from 0 to 1268 nM. (A) Flow cytometry-based quantification of the fraction of fluorescent cells. Error bars reporting binomial standard errors for each technical replicate are smaller than symbols. (B) Bulk relative fluorescence intensity normalized by OD600 (see Materials and Methods), averaged over technical replicates, measured at the plate reader.

In the same experiment, we exposed the biosensor to increasing concentrations of arsenic, from 127 nM to 1.268 μM, to probe its dynamic range and assess its response to contamination levels above the WHO safety limit (133 nM). After exposure for 19 h at 37°C, the biosensor successfully distinguished 127 nM from the zero-arsenic control. Moreover, the response did not reach saturation at the highest concentration (Fig. 5A), suggesting that it may be able to distinguish concentrations higher than those tested here. To test detectability in bulk measurements, we measured bulk fluorescence using the fluorescence plate reader (Fig. 5B). The results show that even a small fraction of recombined cells (∼5% at 127 nM arsenic) can be resolved, thanks to the high fluorescence intensity of individual recombined cells, demonstrating that the difference between the no-arsenic control and 127 nM arsenic remains quantifiable without single-cell resolution and indicating that the recombinase-based biosensor can be used in experimental settings without specialized equipment for anoxic fluorescence measurements.

Different colors show different biological replicates, using the same colors as panel A. Insets compare the As-negative control with the biosensor grown in the absence of arsenic.

Unlike p-ArsRBS2*-syfp2*, the Cre-recombinase based biosensor maintains a memory of arsenite exposure, allowing the user to decouple exposure from measurement. This is an advantage over existing biosensors that use anaerobically compatible fluorescent proteins, which require all aspects of the experiment, including plate reader fluorescence measurements, to be performed inside an anaerobic chamber, and typically suffer from low signal-to-noise ratios due to the reduced brightness of these fluorophores (8,22). We demonstrate this advantage by successfully detecting arsenite exposure in anoxic conditions using the recombinase-based p-ArsRBS*2-cre*[LVA]-loxPP*-syfp2* arsenite biosensor. In a Coy anaerobic chamber, we grew three biological replicates at room temperature in liquid Lysogeny Broth (LB) cultures that had been degassed overnight, using fumarate as an electron acceptor (Fig. 6A). We used LB as a rich nutrient medium to favor *E. coli*’s switch to anaerobic respiration (32) and to increase its growth rate relative to MGP. Cells were exposed for 61 hours to 1 μM arsenite in LB with fumarate and glycerol (to enhance GlpF activity (33)). Cells grown in the same conditions, minus the arsenite, were used as controls. Following exposure, cells were washed in phosphate-buffered saline (PBS) and incubated in MGP in aerobic conditions, at 37 °C, for 24 hours, allowing full maturation of the fluorescent protein and enabling expression in cells that had just recombined the plasmid. Finally, samples were analyzed at the flow-cytometer (Fig. 6B). The fraction of fluorescent cells in samples exposed to arsenite (39% on average) was significantly higher (*P*-value ∼ 10^-17^) than in the control (0.6% on average). We further observed that the subpopulation with only one *loxP* site remained far from saturation after being exposed to a high concentration of arsenite for two days in anoxic conditions. This may indicate that, under anoxic conditions, the biosensor can resolve a wide range of contaminant concentrations or be deployed for long periods in the exposure phase without reaching full recombination of the population.

**Fig. 6.**
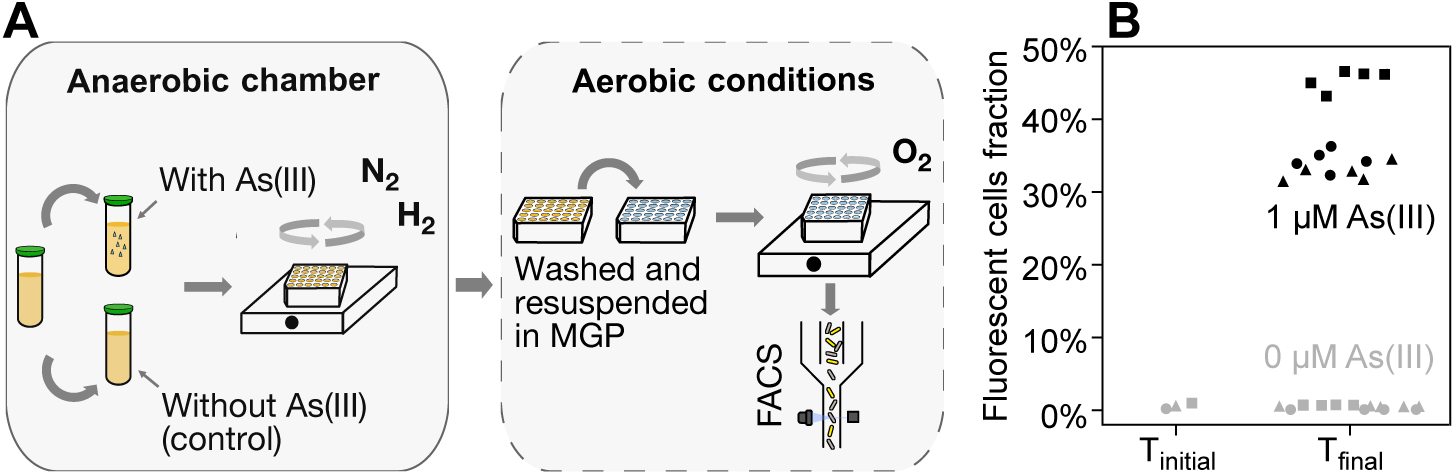
Schematics of the anoxic assay and mean endpoint fluorescence measurements collected at the flow-cytometer for cells carrying p-ArsRBS*2-cre*[LVA]-loxPP*-syfp2*. (A) Schematics of the experimental setup: in the anaerobic chamber (left), cells were grown in well-mixed liquid LB cultures with fumarate and glycerol, re-diluted in the same medium with and without arsenite, and grown in a 96-well plate placed on a microplate shaker for ∼60 h. Samples were then removed from the anaerobic chamber (right), washed to remove arsenite, and resuspended in fresh MGP medium supplemented with dextrose, while lacking arsenite and fumarate. (B) Fraction of fluorescent cells sampled from cultures before transfer into the exposure medium (T_initial_) and at the end of the exposure phase, with or without arsenite (T_final_). Flow-cytometry measurements were performed after cells were resuspended in arsenite-and fumarate-free medium and incubated in aerobic conditions for 24 h, allowing reporter expression to develop. Different symbols represent different biological replicates, different colors represent different arsenite concentrations.

As the resuspended population contains a mixture of fluorescent and non-fluorescent cells, some of which may have only recently transitioned to the recombined state, we examined how the fluorescent fraction changes following transfer from anoxic to oxic conditions. Flow cytometry measurements were collected immediately after sampling from the anaerobic chamber, after 6 hours, and after 24 hours of oxic incubation in arsenite-free medium (Fig. S5). We observed a modest increase in the fraction of fluorescent cells upon exposure to aerobic conditions, with most of the increase occurring within the first 6 hours and smaller changes thereafter. These dynamics are consistent with post-exposure processes such as fluorescent reporter maturation contributing to the observed signal. Notably, applying less-stringent FACS gating criteria yields a constant fraction of fluorescent cells at all time points, indicating that the apparent increase under stringent gating primarily reflects changes in fluorescence intensity rather than changes in population composition. Longer term stability of the reporter after several generations of growth in oxic conditions is discussed below.

Long-term stability of sensing-reporter systems is essential for deploying engineered microbes as reliable environmental biosensors. However, these systems remain vulnerable to failure over successive generations due to mutations or fitness costs associated with the expression of exogenous genetic constructs. To assess the robustness of our designs, we examined the plasmid (p*-cymR-cre*[LVA]-loxPP*-eyfp*) and chromosomal (c*-cre*[LVA]-loxPP*-syfp2*) cumate-sensitive reporters, and the arsenite reporter p-ArsRBS*2-cre*[LVA]-loxPP*-syfp2* for two potential failure modes. First, we wanted to monitor a possible decline in the fraction of fluorescent cells after exposure, which may occur due to fitness costs associated with expressing the fluorescent protein and potentially allow cells that did not perform Cre recombination (with two *lox* sites) to outcompete those that did (with one *lox* site). Alternatively, mutations arising within the latter could silence fluorescent protein expression and sweep through the population. For practical purposes, spontaneous recombination and fluorescent protein expression should be sufficiently rare to avoid false detection of the inducer, although proper controls can help mitigate these issues. Such events compromise the fidelity of the biosensor’s memory and reduce its reliability for long-term quantification of contaminants.

To probe these effects, we measured the fraction of fluorescent cells in the population immediately after transient inducer exposure and tracked this fraction over 50 generations in inducer-free media (Fig.7). We induced the reporters using 2 μM cumate or 500 nM arsenite to generate a subpopulation of fluorescent cells. For the cumate-sensitive reporters, we prepared an additional treatment (5 μM cumate) in which most cells in the population were recombined, allowing us to examine dynamics from both low and high initial recombined fractions. Parallel uninduced cultures were used to monitor background recombination over time. Cells were grown in LB (p*-cymR-cre*[LVA]-loxPP*-eyfp* and c*-cre*[LVA]-loxPP*-syfp2*) or MGP (p-ArsRBS*2-cre*[LVA]-loxPP*-syfp2*), exposed for 12 h, and then diluted every 12 h, with each growth phase corresponding to 13.5 and 12.5 generations, respectively (Materials and Methods). We observed a ∼60% and ∼50% decline in the fluorescent cell fraction after around 50 generations for p*-cymR-cre*[LVA]-loxPP*-eyfp* (Fig. 7A) and p-ArsRBS*2-cre*[LVA]-loxPP*-syfp2* (Fig. 7C), respectively. Conversely, the chromosomal reporter c*-cre*[LVA]-loxPP*-syfp2* (Fig. 7B) showed no reduction in the fluorescent cells fraction, likely reflecting the reduced fitness cost of expressing a single-copy fluorescent protein gene, the absence of plasmid replication burden, and the genetic stability of chromosomal insertions, which are not easily eliminated from the population. After removing the inducer and growing cells in fresh media, we observed an initial increase of the fraction of fluorescent cells in both the chromosomal cumate-sensitive reporter and the recombinase-based arsenite-sensitive reporter. This increase likely reflects recombination that occurred toward the end of the exposure phase, when cells in stationary phase may not yet have had time to express and mature the fluorescent protein.

**Fig. 7.**
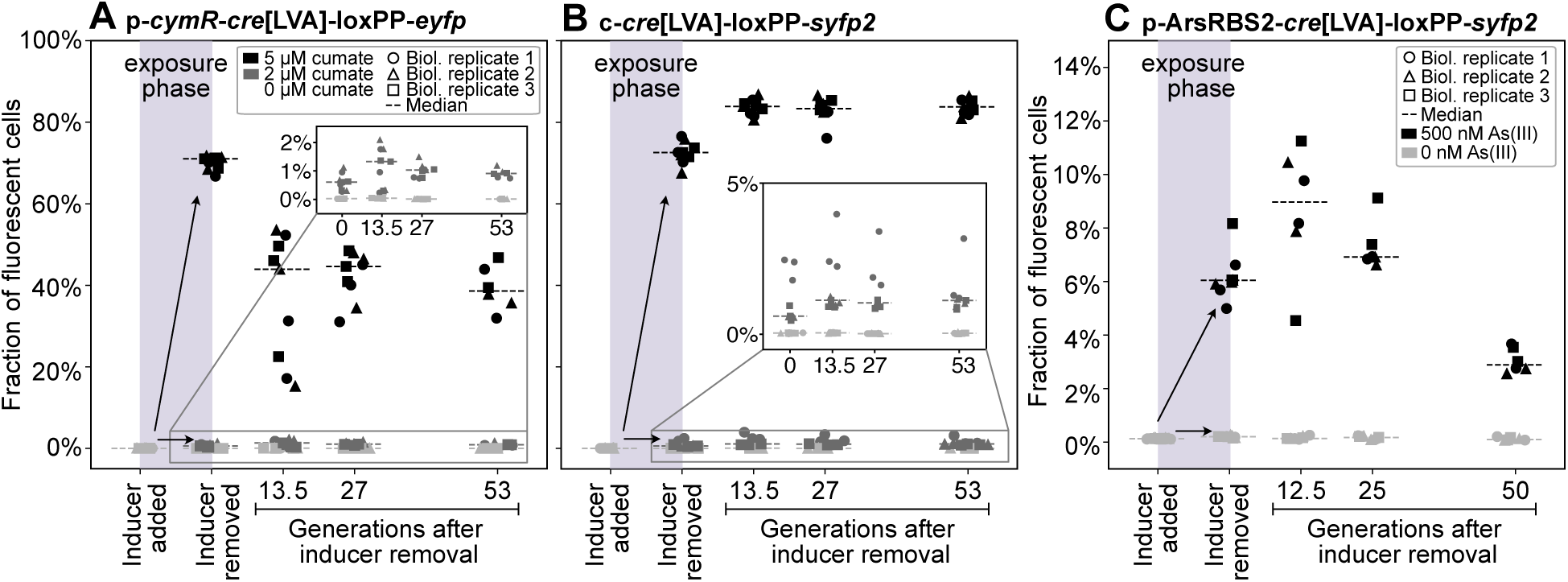
Fraction of fluorescent cells in cultures passaged in inducer-free media after exposure; different panels show data for different Cre reporters: (A) p*-cymR-cre*[LVA]-loxPP*-eyfp*, (B) c-*cre*[LVA]-loxPP-*syfp2*, and (C) p-ArsRBS*2-cre*[LVA]-loxPP*-syfp2*. Cells were measured at the flow-cytometer before the addition of the inducer, after the exposure phase, before washing the cells and removing the inducer, and at several passaging points. Each point is a technical replicate of one of three biological replicates, and it is scattered around the x-axis to aid visualization. Binomial standard errors are smaller than symbols.

## DISCUSSION

To summarize, in this work we explored how different regulatory strategies of a Cre-*lox* system affect the response of cumate-inducible reporters in *E. coli*, on both plasmids and the chromosome. To assess the performance of the engineered bioreporters, we examined their basal expression, sensitivity, titrability, and stability over multiple generations. We observed that the use of an optimized inducible promoter and protein degradation tag effectively limited leakage. Furthermore, we showed that recombination rates could be manipulated by altering the ratio of gene copy numbers of the regulator and transcriptional promoter, and that extremely low recombination rates could be achieved by hindering Cre activity by substituting homologous *loxP* sites with the heterologous *lox511-loxP* pair. Mutated Cre recombinases (34) with reduced efficiency could also be explored to expand the tunability range. Our constructs provide an alternative to optogenetic Cre-*loxP* (16) systems and require only low concentrations of cumate, an inexpensive, non-toxic inducer. Moreover, the dynamic range of induction could potentially be expanded through the addition of salicylic acid, which antagonizes the cumate sensor (23).

To better understand the persistence of genetic memory, we examined reporter stability over multiple generations. As expected, fluorescent protein expression imposes a fitness burden, even when on a low-copy-number plasmid. These fitness costs became detectable after approximately 13 generations following inducer removal. Chromosomal integration improved stability, likely due to reduced gene copy number, lower expression burden, and avoidance of plasmid replication costs. Because our primary goal was to characterize Cre titrability rather than long-term memory stability, we used a constitutively expressed fluorescent protein to facilitate real-time detection of rare recombination events. Future designs could mitigate fitness costs by placing the reporter under inducible control, activated only immediately before measurement.

Recombinases are widely employed in synthetic biology applications for their efficiency, specificity, and ability to encode genetic memory. However, tunable recombinase systems present challenges in prokaryotes, particularly with efficient enzymes like Cre, where tight regulation is needed to prevent unintended genomic changes. Being able to experimentally tune recombinase-mediated recombination rates *in vivo* across a titratable range opens up new possibilities for applications in genetic circuit design (15), cell physiology (14), microbial ecology, and experimental evolution (2) research. For example, low-rate recombination could be used to simulate mutational drift, create stochastic genotypic diversification in structured populations, or trace spatially segregated lineages in microbial colonies and biofilms.

To demonstrate the broad applicability of our circuits, we applied the same design principles to construct a recombinase-based arsenite biosensor. This reporter reliably detected low nanomolar concentrations of arsenite and retained its functionality under anoxic growth conditions. The majority of existing whole-cell arsenite biosensors (29,30) adopt traditional designs comprising fluorescence-based reporter circuits in which transcription of a fluorescent protein is regulated by P*_arsR_*. In contrast, our system uses recombinase-mediated excision to store the exposure signal in DNA, enabling decoupled measurement. We aimed at exploiting the Cre-lox memory to yield a temporally integrated response that depends on exposure duration and population growth. Under this design, even very low inducer concentrations that drive slow recombination become detectable after sufficient exposure, provided that recombination is irreversible, the fluorescent phenotype is genetically stable, and expression imposes minimal fitness cost. This approach prioritizes cumulative dose sensing and dynamic range over rapid switching, enabling robust detection in subsaturation regimes where conventional fast response sensors may be less informative.

Our biosensor’s capability to detect recombination events at low rates and in anoxic conditions could be particularly relevant for monitoring arsenic pollution in real-world environments, such as rice paddy waters, where microbial uptake of dissolved arsenic can be inhibited due to factors including complexation with dissolved organic matter, association with colloidal phases, or repression of arsenic uptake by organic carbon (8,33). Arsenic uptake into microbial cells is a prerequisite for enzymatic arsenic methylation, which produces methylated arsenic species that can represent a significant fraction of the arsenic that accumulates in rice grains (7). Biogeochemical interactions between arsenic, iron, and sulfur that govern arsenic chemical speciation and fate depend strongly on redox conditions, highlighting the need for experimental methods to characterize microbial availability of arsenite under anaerobic conditions where it is most mobile, as found for example in waterlogged paddy soils, wetlands, marine sediments, and deep subsurface environments (7). Although the use of flavin-based fluorescent proteins allows traditional biosensor designs to function in anoxic or anaerobic assays (8,20), the low signal-to-noise ratio of these measurements can significantly reduce the sensitivity and accuracy of detecting low concentrations of analytes. In contrast, recombinase-based biosensors, such as the one developed here, are particularly suitable for these assays thanks to the decoupling of environmental sensing from measurements. This genetic memory brings interesting new functionalities compared to traditional biosensors and can be advantageous for obtaining temporally-integrated measurements in the field or laboratory (13), with cells transitioning to the recombined state at low rates over extended periods of time. In the future, the reporter construct could be further optimized to reduce leakage, for example by disrupting the *arsR* negative feedback loop (35–37), using an engineered version of P*_arsR_*, or integrating the synthetic construct in the chromosome. This would further benefit the inheritance of the inserted sequence across generations without providing a selective antibiotic.

It is important to note that a trade-off of designing the system to operate in a slow recombination regime, suitable for long-term assays, is that very short exposure times or very low analyte concentrations may yield only a small fraction of recombined cells, potentially below the sensitivity of the biosensor. Nevertheless, using high throughput measurements (e.g., via flow cytometry), we were able to reliably detect fractions as low as ∼0.069% when sampling ∼150,000 events, with even lower fractions theoretically accessible at larger sample sizes. Moreover, because recombined cells exhibit a strong fluorescence signal, even small fractions of recombined cells remain detectable in bulk measurements, as evidenced by the results of Fig. 5.

While recombinase-based biosensors are well-established in synthetic biology, their application to environmental sensing remains limited (5). Our findings show that Cre-*lox* systems can be tuned to operate at low recombination rates, maintain stable genetic memory, and function reliably under both aerobic and anoxic conditions. These properties make them well-suited for applications that require temporally integrated sensing, post-exposure readouts, or compatibility with environmentally relevant settings. We note, however, that the host strain used here, *E. coli* DH10B, is an auxotrophic laboratory strain not well suited for growing in environmental settings. While this strain’s characteristics facilitate genetic engineering and laboratory-based analysis of environmental samples, deployment of the constructs developed here would require adopting host strains tailored to the intended application. Further optimization, such as chromosomal integration in environmentally adapted strains, reduction of expression burden, or implementation of alternative signal outputs, could enhance their suitability for field applications.

As synthetic biology continues to expand into environmental and ecological domains, titratable recombinase systems offer a robust and adaptable platform for recording microbial exposure to environmental stimuli and contaminants in dynamic settings.

## MATERIALS AND METHODS

### Strain editing

We assembled and characterized the cumate reporter in the Marionette-Clo *E. coli* strain (23), which has DH10B background. On plasmids, *eyfp* transcription is controlled by the constitutive, medium-strength promoter P_W4_ (16). In the chromosomal reporter, *syfp2* (38) transcription is regulated by the high-strength constitutive promoter P*_rpsL_*. P*_rpsL_-syfp2* was amplified from strain eDE31, which was a gift from Johan Paulsson. Cre was a gift from Niels Geijsen (Addgene plasmid #62730) (39). The terminators *ECK120033736 and L3S2P11* and *lox511* site were synthesized by Twist Bioscience (Table S5). P_W4_ and *loxP* were synthetized by Twist Bioscience, sequences are from Sheets et al. (16). Plasmids were constructed using FastCloning (40) or Gibson assembly (41). Oligonucleotides used are reported in Table S5. Chromosomal insertions were achieved via the mTn7 (MCS MS26) mobile element, inserting at *attTn7*, downstream of *glmS*, following the protocol in Sibley & Raleigh (42). Plasmids were isolated with the Omega Bio-Tek E.Z.N.A. Plasmid DNA Mini Kit I (Q - spin) and sequenced via Oxford Nanopore long-read sequencing. Chromosomal insertions were amplified with primers oAG70/oEG103 (Table S5) surrounding *attTn7* and sequenced via Oxford Nanopore long-read sequencing. After substituting *mCherry* in pMP01 with the *cre*[LVA]-loxPP*-syfp2* construct, we transferred the reporter sequence on a vector with low copy number (pAJM.717, with p15A origin of replication). This decreased the leakage of the reporter. The p-ArsRBS*2-cre*[LVA]-loxPP*-syfp2* and p-ArsRBS2*-syfp2* constructs were assembled via FastCloning, except for the last step, which consisted of swapping the backbone with the low copy number vector and was performed via enzymatic restriction cloning.

### Media and chemical inducers

Characterization of the cumate-inducible constructs was performed in Lysogeny Broth (LB) medium with the appropriate selective antibiotic, whereas the arsenite-inducible constructs characterization in aerobic conditions was done in MOPS for growth medium (MGP), following the recipe used in Pothier *et al.* (30), supplemented with 50 μg/μL ampicillin sodium salt. Super Optimal broth with Catabolite repression (SOC) medium and plates made with Lysogeny Broth (LB) medium with 1.5% agar and 50 μg/μL kanamycin sulfate were used for cloning and strain maintenance. Unless specified otherwise, cells were grown in LB with the appropriate selective antibiotic. To induce *cre* transcription, we added appropriate amounts of a stock solution of 100 mM cuminic acid (4-isopropylbenzoic acid) ≥ 96% purity (Sigma-Aldrich 268402) in 100% ethanol to the medium.

### Cumate-sensitive reporter response characterization

Plate reader assays: Overnights of strains of interest were grown from fresh single colonies on agar at 37 °C in 1 mL of LB in a roller drum. The following day, cultures were diluted 1:200 in glass culturing tubes and incubated at 37 °C on the roller drum for 2 hours. Afterwards, cells were further diluted 1:100 in pre-warmed LB with antibiotic and inducer and, for each sample and treatment, a total of 1.5 mL was distributed in equal volumes of 125 μL into 12 wells of a sterile, flat-bottom, non-treated, 96-well tissue culture plate with lid (VWR 10861-562). Three technical replicates were used for each treatment condition and time-step recovery. The samples were distributed on the plate in a random configuration. Cells were then grown in the plate reader at 37 °C with continuous double orbital shaking at 807 cpm. Strains were induced with a range of cumate concentrations ranging from 0 μM to the maximum concentration of cumate as recommended in Meyer *et al.* (23), i.e. 100 μM. Some biological replicates showed a fitness cost when grown at this maximum concentration (Table S2). OD600 and fluorescence (excitation at 501 nm and emission at 533 nm) were measured every 15 minutes over 21 hours of incubation. OD600 and fluorescence data of 3 wells inoculated with only medium and no cells were averaged at each timestep and used as a measure of background expression. At four time points, after 1 h, 2 h 30 min, 5 h and 21 h from the beginning of the assay, the plate was shortly removed from the plate reader, between 2 and 5 μL were transferred from each well into a volume ranging from 125 to 198 μL of Tris-HCl pH 7.5 in a 96-well U bottom plate to perform flow-cytometry measurements (10,000 single cells recorded per sample).

The relative fluorescence intensity output (Figs. 2 and 5B) was computed as follows:

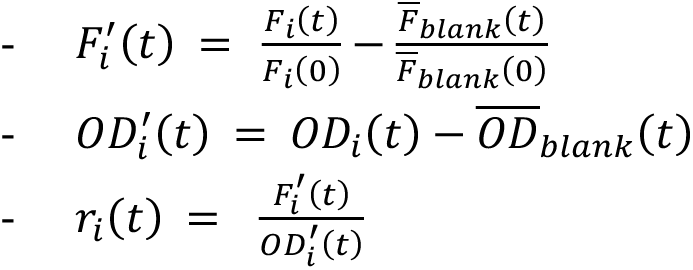

where 𝐹_i_ (𝑡) is the fluorescence intensity read for well 𝑖 at time point 𝑡, 𝐹_i_(0) is the fluorescence intensity read at the start of the experiment, and 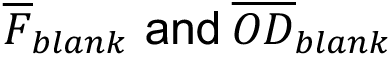 are the fluorescence intensity and OD600 values averaged over three wells containing medium. 𝑟_i_(𝑡) is the value plotted in Figs. 2 and 5B.

Flow-cytometry and colony PCR: Flow-cytometry was performed by collecting forward and side scattering (FSC/SSC), as well as fluorescence intensity with excitation at 488 nm and emission at 533/30nm (FL1). We performed data analysis using the Python package FlowCytometryTools (44). In the biological replicates experiment shown in Fig. S1(D-F), three technical replicates for each treatment at different times were pooled and used as template for colony PCRs to amplify the DNA region enclosing the two *loxP* sites. After qualitatively comparing the bands length difference, the DNA was purified from the PCR product with the Omega Bio-Tek E.Z.N.A. Cycle Pure Kit (V-Spin) and Sanger sequenced. The generated chromatograms were inspected to confirm the presence of the recombined and non-recombined variants. In all experiments, the parent strain Marionette-Clo was used as a negative control.

Recombination rates estimation: To estimate recombination rates for p*-cymR-cre*[LVA]-loxPP*-eyfp* and c*-cre*[LVA]*-*loxPP-*syfp2* as a function of cumate concentration, we fitted a logistic curve to flow cytometry data (Fig. 3):

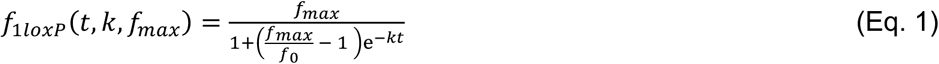

where:

-𝑓_0_ is the experimentally measured fraction of fluorescent cells at time 0,
-𝑓*_max_* is a fitting parameter, bounded between the maximum measured fraction and 1,
-𝑘 is a fitting parameter representing the recombination rate (h^-1^),
-𝑡 is time in hours.

At low cumate concentrations, the parameters of the logistic growth model are not identifiable because a time-invariant fraction of fluorescent cells can arise either by setting 𝑓*_max_* = 0 (for any 𝑘), or setting 𝑘 = 0 (for any 𝑓*_max_*) in Eq. 1. Furthermore, if 𝑓 ≪ 𝑓*_max_* throughout the time course, 𝑓*_max_* cannot be determined. Thus, for each technical replicate, we fit both the logistic Eq. 1 and its asymptotic expansion in the small parameter 𝑓/𝑓*_max_*, whose leading term is the exponential growth 𝑓(𝑡, 𝑘) = 𝑓_0_𝑒*^kt^*, which has one fewer fitting parameter compared to Eq. 1. Accordingly, for each technical replicate we fit both the full logistic model (maximum likelihood fit of Eq. 1) and its early-growth exponential approximation, and we select the estimate of 𝑘 from the model with the lowest corrected Akaike Information Criterion (AICc) value. A non-linear fit was performed for the exponential model, while Markov chain Monte Carlo was used for the logistic model fit. Only the exponential fit was performed for replicates with no supplemented cumate. Figs. 3 and S1 and Table S3 report the average across technical replicates of such best fits.

### Arsenite-sensitive reporter response characterization and analysis Oxic conditions

For the experiments of Fig. 4, a stock solution of 7 mM of arsenite was prepared by dissolving sodium arsenite (LabChem, Zelienople, PA) in Milli-Q water, followed by 0.22 μm syringe-filtration. The preparation was conducted in a Coy anaerobic chamber (98% N₂ and 2% H₂), and the stock solution was stored anaerobically at 4 °C protected from light after its concentration was verified using ICP-MS (Agilent 7800). From this stock, we collected 15 μL diluted in a total volume of 15 mL of sterile water in a 15 mL falcon tube. Experiments were carried out in the same week the 70 μM arsenite stock was prepared, to minimize the solution exposure to air and the risk of oxidation into As(V). Yoon *et al.* (45) verified the absence of arsenite speciation in their biosensor experiments via HPLC-ICP-MS analysis, with sampling conducted during the first 10 h of their experiment, under conditions similar to our experiments. For the experiment of Fig. 5, a ∼1 mM stock solution of arsenite was diluted 10-fold and filter sterilized in oxic conditions, and stored at room temperature protected from light, within one week from the experiment. In parallel, the arsenic concentration of the working solution was quantified by ICP-MS.

The host strain used for the arsenite biosensors is *E. coli* NEB 10-beta, which has DH10B background. Strains used for the arsenite-sensitive reporter assay were streaked from the cryostock on an LB agar plate with 2% dextrose, 50 μg/mL kanamycin sulfate, and 50 μg/mL ampicillin sodium salt and incubated at 37 °C overnight. At least three colonies per strain were picked and inoculated into 2 mL-deep 96 well plates (VWR 76397-576) with 1 mL of MGP supplemented with 50 μg/mL ampicillin sodium salt and 100 μL of 20% dextrose. The plate was covered with a rayon film and lid and placed on an orbital shaker rotating at 400 rpm in a 37 °C incubator. The following day, samples were measured at the flow cytometer. Dextrose helped reduce Cre expression leakage, probably due its inhibitory action on the glycerol facilitator transporter protein (GlpF), which mediates the uptake of arsenite by the cell (33,46). The biological replicates with the least leakage were selected for the assay. Cells were washed in MGP to remove dextrose and diluted 10,000 fold in MGP with ampicillin and different concentrations of arsenite. Technical replicates were inoculated in 2 mL-deep 96-well plates and kept in the incubator with 400 rpm orbital shaking until they reached stationary phase. Samples were measured at the flow cytometer after 17 hours of incubation (18 hours for the third biological replicate) by sampling between 10 and 12 μL of liquid, which were then diluted in 50 mM Tris-HCl pH 7.5. For each well, we collected 150,000 single-cell measurements. The binary response of the recombinase-based biosensor was measured at the single-cell level by measuring the fraction of cells contained within a gate in the FSC-A vs FL1-A that was drawn by comparing cells grown in the absence of As(III) to fully recombined cells.

To assess statistical significance (*P*-value < 0.05) of the sensors’ sensitivity, within the same biological replicate, we performed two-sided Welch’s *t*-tests between treatments at successive inducer concentrations (i.e., 0 nM with 15 nM, 15 nM with 30 nM, etc). The *P*-values obtained from the different experiments (biological replicates) for the same arsenite concentration were then combined following Fisher’s method. The corresponding *P*-values are reported in Table S4. Because the third biological replicate had a slightly longer duration than the first two, we also report in Table S4 combined *P*-values for the first two experiments only, excluding the third one.

### Anoxic conditions

Experiments were conducted at room temperature in a Coy chamber with 97.7% N_2_ and H_2_ 2.3% and O_2_ < 20 ppm. p-ArsRBS2*-cre*[LVA]*-*loxPP*-syfp2* was streaked on plates prepared by mixing in aerobic conditions one part of 4% agar with one part of MGP medium with 10mM sodium fumarate and grown in low oxygen conditions (O_2_ around 1000 ppm). Plates were supplemented with 2% dextrose (to repress GlpF expression), 50 μg/mL ampicillin sodium salt, and 50 μg/mL kanamycin sulfate. Growth and exposure were performed in liquid LB medium supplemented with 80 mM fumarate, 0.001% resazurin and 50 μg/mL ampicillin sodium salt. Additionally, the exposure medium was supplemented with 0.4% v/v glycerol. Media were sterilized by autoclaving, allowed to cool below 60 °C before adding ampicillin, and then transferred into the anaerobic chamber. Bottle lids were left loosely closed overnight to promote degassing, and a rayon film was placed over the opening to prevent contamination. Degassing was confirmed using a resazurin dye test, which also served as a visual indicator of anoxic conditions during cell growth (Fig. S4). Three isolated colonies were picked and inoculated into 2 mL of medium in 10 mL Falcon tubes and placed on a shaker rotating at 160 rpm. After 6 days of growth in liquid LB, 10 μL/mL of the cultures were transferred to a 2 mL-deep 96-well plate, containing 1 mL per well of exposure medium, with or without 1 μM arsenite. We prepared five technical replicates for each of the three biological replicates. The plate was covered with rayon film and a lid and incubated on a shaker set at 150 rpm. A 50 µM arsenite stock solution was prepared under an air atmosphere and sterile-filtered into an autoclaved serum bottle. The bottle was then capped, crimped, and its headspace was purged with argon for 15 minutes. Cells were exposed to the treatment conditions for 61 hours. At the end of the exposure period, the cultures were removed from the anaerobic chamber, immediately washed in PBS, diluted 1:100, and transferred to MGP supplemented with 2% dextrose for aerobic culturing. The 2 mL-deep 96-well plate was incubated at 37 °C on a shaker rotating at 400 rpm for 24 hours to allow full maturation of the fluorescent protein. For the initial time point t_0_, 100 μL of the LB culture (prior to inoculation) was taken out of the chamber, resuspended in MGP without fumarate (10 uL in 1 mL total volume), and grown under the same conditions for 24 hours. After incubation, cells were diluted 10-fold in PBS supplemented with 15 μg/mL propidium iodide and analyzed by flow cytometry to assess viability. No differences in viability were observed in the presence and absence of arsenite (*P*-value 0.41, *Fig*. S5).

### Reporters’ stability assay

The plasmid and chromosomal cumate reporters were streaked on LB plates supplemented with 50 μg/mL kanamycin sulfate. p-ArsRBS2*-cre*[LVA]*-*loxPP*-syfp2* was streaked on an LB plate with 50 μg/mL of kanamycin sulfate, 50 μg/mL of ampicillin sodium salt, and 2% dextrose. The day after streaking, three colonies from each plate were inoculated into 1 mL of medium (for p-ArsRBS2*-cre*[LVA]*-*loxPP*-syfp2* MGP with kanamycin and 2% dextrose). Overnights were diluted 10,000-fold into 1 mL of medium without inducers and distributed in a 2 mL-deep 96-well plate, covered with rayon film and placed on a rotating roller drum. After 12 hours of incubation, cells were diluted in fresh media with inducers: 0, 2 or 5 μM of cumate, and 0 or 500 nM of arsenite. From here onwards, dextrose was absent from MGP. After 12 hours, cells were diluted 100-fold in PBS, followed by another 100-fold dilution in medium without inducers. Dilution and propagation into fresh media were carried out every 12 hours for five cycles. Every 24 hours, cells from the intermediate 100-fold dilution in PBS were measured at the flow cytometer. We measured 200,000 cells per sample, or up to 140 uL, whichever occurred first. Following this protocol, p-*cymR-cre*[LVA]-loxPP-*eyfp* and c*-cre*[LVA]-loxPP*-syfp2* went through ∼53 generations after exposure. In MGP, p-ArsRBS2-*cre*[LVA]*-*loxPP*-syfp2* grows slower than the other bioreporters in LB and does not reach carrying capacity after 12 hours. Therefore, the number of cell generations per cycle was computed as log_2_ 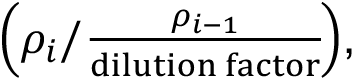 where 𝜌 is the total number of cells counted at the flow-cytometer, divided by the volume at the 𝑖-th cycle.

## Author Contributions

E.G. and A.G. conceived the study. E.G. performed the experiments and data analysis. A.G. supervised the experiments and data analysis. H.Y. and M.C.R. provided the arsenite stock solution, gave technical and conceptual advice for the arsenite biosensor assays, and provided support for anoxic experiments. E.G. and A.G. wrote the manuscript. H.Y. and M.C.R. edited the manuscript.

## ACKNOWLEDGEMENTS

We thank Andrew Murray for insightful comments on the manuscript and Katie Randolph for help with the experiments. We thank Michael A. P. Vega for his support with the preparation of degassed media for the anaerobic chamber, the preparation of sodium arsenite and ICP-MS arsenic concentration assays. We thank the Cornell Statistical Consulting unit and the Cornell. We thank the Genomics Facility (RRID:SCR_021727) of the Biotechnology Resource Center of Cornell Institute of Biotechnology for their help with sequencing experiments. AG acknowledges support by the National Institute of General Medical Sciences of the National Institutes of Health, United States of America under award number 1R35GM147493 and by the Human Frontier Science Program award no. RGEC28/2023. MCR acknowledges support from NSF grant 1905175.

## DATA AVAILABILITY

All plate reader and flow cytometer data, and corresponding data analysis scripts are available on Zenodo https://doi.org/10.5281/zenodo.18705642, along with sequencing data used for strain and plasmid validation.

Plasmids p*-cymR-cre*[LVA]-loxPP*-eyfp*, p-ArsRBS*2-cre*[LVA]-loxPP*-syfp2*, strain c*-cre*[LVA]*-*loxPP-*syfp2* were deposited on Addgene.

## SUPPLEMENTAL MATERIALS

Supplemental materials include supplementary tables reporting strains’ growth rates measured in the experiments carried out to characterize the cumate-sensitive reporter population, *P*-values of all statistical tests performed, the main oligonucleotides used in this study, and additional notes on data analyses.

