## Supplemental Materials for "Tunable Low-Rate Genomic Recombination with Cre-*lox* in *Escherichia coli*: A Versatile Tool for Anoxic Environmental Biosensing and Synthetic Biology"

Supplemental materials include supplementary tables reporting strains' growth rates measured in the experiments carried out to characterize the cumate-sensitive reporter population, *p*-values of all statistical tests performed, the main oligonucleotides used in this study, and additional notes on data analyses.

### Table S1 Recently developed biosensors.

The whole-cell biosensors reported in the table for comparison with the system constructed in this work present novel types of output signals and/or high sensitivity. We specify statistical method used to determine the limit of detection of the sensor and the linear dose-response range, when specified.

| Strain and reporter name | Ref | Output | Sensitivity / Limit of detection [nM] | Detection range [nM] | Incubation time and readout |
|---|---|---|---|---|---|
| *E. coli DH5α* pJAMA-arsR-ABS | (1) | LuxAB (Luminescence) | 50 (2-fold signal increase from blank) | 50 – 800 (linear response 50-500) | 1 hour - luminometer |
| *E. coli* TOP10 arsR1-$P_{ars}$-GFP | (2) | GFP (Fluorescence) | 10 (student t-test p-value < 0.05) | 30 – 100 (linear range) | 4 hours – microplate reader |
| *E. coli DH5α* ParsG-ABS-8 | (3) | GFP (Fluorescence) | 10 (2-fold signal increase from blank) | 75 – 4000 | 4 hours - microplate reader |
| *E. coli NEB5- α* pAH1T-$P_{ars}$ -PpFBFP | (4) | PpFBFP (Fluorescence) | 250 | 250 – 5000 | 10 hours - anaerobic, microplate reader |
| *E. coli* TOP10 TOP10/pK12-DV | (5) | Deoxyviolacein (Pigment) | 1 (blank + 1.65 SD) | 45 - 586 | 4 hours - absorbance |
| *E. coli* TOP10 TOP10/pnK12-ABS-ind | (6) | Indigoidine (Pigment) | 45 (blank + 1.65 SD) | 45 – 18750 (linear) | 5 hours - microplate reader |
| *E. coli DH5α* pPR-*arsR*-ABS-RBS-*lacZ* | (7) | LacZ + X-gal as substrate (Colorimetric) | 2.7 | 2.7 – 133 (linear response 2.7-80) | 8 hours - microplate reader |
| *E. coli DH5α* FRED-As | (8) | LacZ + substrate 4-aminophenol (Electrochemical) | 30 (MDL) | 30-1064 (linear 30-266) | 1 hour - single-use cartridge |
| *E. coli NEB10-beta* p-ArsRBS2-*cre*[LVA]-loxPP-*syfp2* | This study | SYFP2 (Fluorescence) | 15 (student t-test p-value < 0.05) | 15 – 1268 | 17 – 20 hours incubation – flow-cytometry |

### Table S2 Growth rates measured for each experiment shown in Fig. 2.

Growth rates computed for each technical replicate by fitting a linear model to the plate reader $\ln(OD600)$ data versus time, during the exponential growth phase. The blank medium OD was subtracted from the data prior to fitting. Fits were performed using data points that fell within the same range of OD values ([0.03,0.1]), for each strain. We report the average growth rate across three technical replicates, for each of the three biological replicates, organized as median value and minimum/maximum range in square brackets. Welch's t-test was computed for each strain and concentration compared with the condition without cumate for each treatment. We report the combined *p*-value across the biological replicates via Fisher's method, corrected for multiple testing using the Benjamini–Hochberg false discovery rate procedure.

| **Strain and corresponding construct** | **Cumate concentration** | **Growth rate median [min,max] [$h^{-1}$]** | **Fisher-combined *p*-value** | **BH adjusted *p*-value** |
|---|---|---|---|---|
| bEG36 (*p-cymR-cre*[LVA]-loxPP-*eyfp*) | 0 µM | 0.94 [0.92,1.03] | | |
| bEG36 (*p-cymR-cre*[LVA]-loxPP-*eyfp*) | 3 µM | 0.93 [0.92,1.05] | 0.66 | 0.76 |
| bEG36 (*p-cymR-cre*[LVA]-loxPP-*eyfp*) | 5 µM | 0.94 [0.90,1.03] | 0.76 | 0.76 |
| bEG36 (*p-cymR-cre*[LVA]-loxPP-*eyfp*) | 8 µM | 0.925 [0.914,1.05] | 0.59 | 0.76 |
| bEG36 (*p-cymR-cre*[LVA]-loxPP-*eyfp*) | 10 µM | 0.90 [0.88,1.041] | 0.64 | 0.76 |
| bEG39 (p-*cre*[LVA]-loxPP-*eyfp*) | 0 µM | 0.93 [0.93, 0.97] | | |
| bEG39 (p-*cre*[LVA]-loxPP-*eyfp*) | 2 µM | 0.95 [0.92,1.0] | 0.76 | 0.81 |
| bEG39 (p-*cre*[LVA]-loxPP-*eyfp*) | 4 µM | 0.94 [0.89, 0.97] | 0.81 | 0.81 |
| bEG39 (p-*cre*[LVA]-loxPP-*eyfp*) | 10 µM | 0.93 [0.90, 1.01] | 0.63 | 0.81 |
| bAG56 (p-*cymR-cre*[LVA]-lox511P-*eyfp*) | 0 µM | 0.96 [0.89,1.05] | | |
| bAG56 (p-*cymR-cre*[LVA]-lox511P-*eyfp*) | 10 µM | 0.97 [0.95,1.05] | 0.71 | 0.71 |
| bAG56 (p-*cymR-cre*[LVA]-lox511P-*eyfp*) | 50 µM | 0.93 [0.89,1.00] | 0.25 | 0.38 |
| bAG56 (p-*cymR-cre*[LVA]-lox511P-*eyfp*) | 100 µM | 0.84 [0.84,0.98] | 0.03* | 0.08 |
| bEG57 (c-*cre*[LVA]-loxPP-*syfp2*) | 0 µM | 0.92 [0.97,1.03] | | |
| bEG57 (c-*cre*[LVA]-loxPP-*syfp2*) | 3 µM | 0.94 [0.93,1.01] | 0.53 | 0.90 |
| bEG57 (c-*cre*[LVA]-loxPP-*syfp2*) | 5 µM | 0.96 [0.91,1.038] | 0.70 | 0.90 |
| bEG57 (c-*cre*[LVA]-loxPP-*syfp2*) | 8 µM | 0.94 [0.93,1.00] | 0.36 | 0.90 |
| bEG57 (c-*cre*[LVA]-loxPP-*syfp2*) | 10 µM | 0.96 [0.92,1.04] | 0.90 | 0.90 |

**Table S3 Cumate reporter recombination rates estimates**

Estimated mean recombination rates across technical replicates and associated standard deviations. They were computed for different treatments by fitting the cytometry data with a logistic curve (Eq. S1) as described in Materials and Methods.

| | **Plasmid reporter recombination rates ($h^{-1}$)** | | |
|---|---|---|---|
| **[Cumate]** | **Exp. 1** | **Exp. 2** | **Exp. 3** |
| 0 µM | -0.16 ± 0.06 | -0.3 ± 0.2 | 0.38 ± 0.03 |
| 2 µM | | | 1.3 ± 0.6 |
| 3 µM | 0.36 ± 0.02 | 0.33 ± 0.07 | 1.33 ± 0.06 |
| 5 µM | 0.780 ± 0.005 | 0.766 ± 0.005 | 1.60 ± 0.05 |
| 10 µM | 1.82 ± 0.04 | 1.85 ± 0.02 | 4.7 ± 0.1 |
| | **Chromosomal reporter recombination rates ($h^{-1}$)** | | |
| **[Cumate]** | **Exp. 1** | **Exp. 2** | **Exp. 3** |
| 0 µM | -0.17 ± 0.06 | -0.06 ± 0.02 | -0.21 ± 0.03 |
| 4 µM | | | 0.3 ± 0.3 |
| 6 µM | 0.59 ± 0.02 | 0.654 ± 0.008 | 1.22 ± 0.02 |
| 8 µM | 0.87 ± 0.02 | 0.93 ± 0.02 | 2.106± 0.02 |
| 10 µM | 1.16 ± 0.08 | 1.26 ± 0.04 | 3.2 ± 0.2 |

**Table S4 *P*-values for the arsenite biosensors sensitivity assay (Fig. 4)**

Individual Welch's t-test *p*-values comparing treatments within biological replicates, as well as the combined *p*-value obtained with Fisher's method and corrected for multiple testing (Benjamini–Hochberg FDR method). We also provide the adjusted combined p-value excluding replicate 3 which had a longer exposure duration, showing that exclusion of the third replicate does not alter the statistical test outcome (significant if *P*-value < 0.05).

- Aerobic assay

| | **Recombinase-based biosensor p-ArsRBS*2-cre*[LVA]-loxPP-*syfp2*** | | | | |
|---|---|---|---|---|---|
| **Comparison** | **Exp. 1** | **Exp. 2** | **Exp. 3** | **Fisher-combined & BH-adjusted (1-3)** | **Fisher-combined & BH-adjusted (1-2)** |
| 0 - 15 nM | 0.24 | $2.4 \times 10^{-2}$ | $1.1 \times 10^{-4}$ | $7.4 \times 10^{-5}$ | 0.035 |
| 15 - 30 nM | $2.0 \times 10^{-3}$ | $6 \times 10^{-5}$ | $4.1 \times 10^{-3}$ | $6.0 \times 10^{-8}$ | $4.08 \times 10^{-6}$ |
| 30 - 45 nM | $5.3 \times 10^{-4}$ | $4.9 \times 10^{-4}$ | $6.7 \times 10^{-4}$ | $7.7 \times 10^{-8}$ | $5.6 \times 10^{-6}$ |
| 45 - 60 nM | $1.1 \times 10^{-4}$ | $4.8 \times 10^{-4}$ | $4.2 \times 10^{-3}$ | $7.7 \times 10^{-8}$ | $3.6 \times 10^{-6}$ |

| | Transcription-based biosensor p-ArsRBS*2*-*syfp2* | | | | |
|---|---|---|---|---|---|
| **Comparison** | **Exp. 1** | **Exp. 2** | **Exp. 3** | **Fisher-Combined & BH-adjusted (1-3)** | **Fisher-Combined & BH-adjusted (1-2)** |
| 0 - 15 nM | $7.0 \times 10^{-2}$ | $1.7 \times 10^{-4}$ | $2.3 \times 10^{-2}$ | $1.4 \times 10^{-4}$ | $3.0 \times 10^{-4}$ |
| 15 - 30 nM | $1.3 \times 10^{-2}$ | 0.13 | $8.8 \times 10^{-2}$ | $9.2 \times 10^{-3}$ | 0.017 |
| 30 - 45 nM | $3.0 \times 10^{-4}$ | $7.0 \times 10^{-3}$ | 0.38 | $1.8 \times 10^{-4}$ | $1.2 \times 10^{-4}$ |
| 45 - 60 nM | $4.4 \times 10^{-2}$ | 0.37 | 0.38 | 0.119 | 0.08 |

- Anoxic assay

| | Recombinase-based biosensor (p-ArsRBS*2*-*cre*[LVA]-loxPP-*syfp2*) | | | |
|---|---|---|---|---|
| **Comparison** | **Exp. 1** | **Exp. 2** | **Exp. 3** | **Combined** |
| 0 - 1 µM | $8.03 \times 10^{-7}$ | $4.8 \times 10^{-7}$ | $2.2 \times 10^{-7}$ | $8.52 \times 10^{-17}$ |

**Table S5 Oligonucleotides used in this study.**

| Oligo | Sequence $5' \rightarrow 3'$ | Description |
|---|---|---|
| oAG90 | atcagaccggaaagcacatc | Forward primer for cre-*lox* construct insertion into pAJM.657 |
| oAG91 | AGGACGAAACAGCCTCTAC | Reverse primer for cre-*lox* construct insertion into pAJM.657 |
| oAG92 | tgctttccggtctgatgaAAGAGGGGAAATACTAGA | Forward primer to amplify *cre* from Addgene plasmid #62730 |
| oAG93 | tcgtcattgatcaaagcTTAAGCTACTAAAGCGTAGTTTT | Reverse primer to amplify *cre* from Addgene plasmid #62730 |
| oAG94 | acgctttagtagcttaaGCTTTGATCAATGACGAACAATAAGG | Forward primer to amplify $P_{W4}$-*lox511*-TT-*loxP* synthesized DNA for insertion in pAJM.657 |
| oAG95 | gaggctgtttcgtcctTCCTTCTTAAAATAACTTCGTATAATGTATGC | Reverse primer to amplify $P_{W4}$-*lox511*-TT-*loxP* synthesized DNA for insertion in pAJM.657 |
| oAG129 | ccttttcggcatcgccctaaaattcggAGAGGGACACGGCGAAATAA | Forward primer used for intermediate plasmid reporter optimization |
| oAG130 | agggcgatgccgaaaaggtgtcaagaaatataCCAGAAACAAAAAAAGGCCGCG | Reverse primer to substitute $P_{W4}$ with $P_{rpsL}$ in pEG36 |
| oAG131 | aaagaggagaaatactagATGGTGAGCAAGGGCGAGGA | Forward primer to introduce the BBa_B0034 RBS upstream of the fluorescent protein in pEG36 |
| oAG132 | ctagtatttctcctctttGAGGCTGTTTCGTCCTTCCTTCT | Reverse primer used for intermediate plasmid reporter optimization |
| oEG96 | gattcaggaatacagaGATAAGGATCCTAATTGGT | Forward primer to delete $cymR^{AM}$ in pEG36/pAG56 |
| oEG97 | tctgtattcctgaatcCATTGGACCAAAACGAA | Reverse primer to delete $cymR^{AM}$ in pEG36/pAG56 |
| oEG98 | tatagcatACATTATACGAAGTTATCAAACGCATG | Forward primer to replace *lox511* with *loxP* in pAG56 |
| oEG99 | tataatgtatgCTATACGAAGTTATTTCGCCGTGT | Reverse primer to replace *lox511* with *loxP* in pAG56 |

| | | |
|---|---|---|
| oEG89 | tattttaattaaTTAGAAAAACTCATCGAGCATCA | Forward primer to amplify reporter and add PacI restriction site to pEG55 |
| oEG123 | attactcgagAGTGCCAACATAGTAAG | Reverse primer to amplify reporter and add XhoI restriction site to pEG55 |
| oEG48 | CCTGATTTATGGCGCGAAAG | Forward primer in *cre* for colony PCR |
| oEG102 | CTGAACTTGTGGCCGTTTAC | Reverse primer in *eyfp* for colony PCR |
| oEG116 | CCGTATGTTGCATCACCTTC | Reverse primer in *syfp2* for colony PCR |
| oAG70 | AATCTGTAACGTTCCGGGTTC | Forward primer downstream of *attTn7* to verify integration in the chromosome via colony PCR |
| oEG103 | CGGTGGTACGCATAACTTTC | Reverse primer downstream of *attTn7* to verify integration in the chromosome via colony PCR |
| oEG112 | agaaatactagATGAGTAAAGGAGAA | Forward primer to amplify *syfp2* from bAG10 chromosome |
| oEG113 | tatttgctcaGATTTGTCCTACTCA | Forward primer to amplify *syfp2* and its terminator from bAG10 chromosome |
| oEG114 | aggacaaatcTGAGCAAATATTTTATCTGAGG | Forward primer used for plasmid reporter fluorescent output optimization |
| oEG115 | ctttactCATCTAGTATTTCTCCTCTTTG | Reverse primer to replace *eyfp* with *syfp2* |
| oEG150 | ttttctccttcattaCAGTCACGACGTTGT | Forward primer to remove pRO1600 *rep* and *oriV* from pMP01 |
| oEG151 | taatgaaggagaaaaGTCAGGTGGCACTTT | Reverse primer to remove pRO1600 *rep* and *oriV* from pMP01 |
| oEG144 | gagtaggacaaatcGTTTGCGTATTGGGC | Forward primer on pMP01 to replace *mCherry* with *syfp2* or the *cre* – *loxP* system |
| oEG145 | tccggtgacagctaGTGATGGTGATGGTG | Reverse primer on pMP01 to replace *mCherry* with the *cre* – *loxP* system |
| oEG146 | accatcaccatcacTAGCTGTCACCGGAT | Forward primer to amplify *cre*[LVA] – $P_{rpsL}$ – loxPP– *syfp2* from pEG55 and insert them on pMP01 |
| oEG147 | CCCAATACGCAAACGATTTGTCCTACTCA | Reverse primer to amplify *cre*[LVA] – $P_{rpsL}$ – loxPP– *syfp2* from pEG55 and insert them on pMP01 |
| oEG148 | accatcaccatcacGAAACAGCCTCAAAG | Forward primer on pEG55 to amplify *syfp2* |
| oEG149 | tttgaggctgtttcGTGATGGTGATGGTG | Reverse primer on pMP01 to replace *mCherry* with *syfp2* |
| oEG172 | ggcgttaattaaACGAAAACTCACGTTAAG | Forward primer to add PacI restriction site to the ArsRBS2 reporter system and transfer it on a low-copy number backbone |
| oEG173 | atttctcgagCAATACGCAAACGATTTGTC | Reverse primer to add XhoI restriction site to the ArsRBS2 reporter system and transfer it on a low-copy number backbone |
| oEG182 | aaatctcgagCTCGGTACCAAATTCCAGAA | Forward primer to add XhoI restriction site to pAJM.717 to insert the arsenite reporter system |
| oEG183 | gcgattaattaaGTTAAGCGGGAGACCAGAA | Reverse primer to add PacI restriction site to pAJM.717 to insert the arsenite reporter system |
| oEG320 | ttccccgaAAGCTTTGATCAATGACGAACAATAAG | FWD primer on pEG69 to build As-negative control |

| | | |
|---|---|---|
| oEG321 | tgatcaaagcTTTCGGGGAAATGTGCGC | REV primer on pEG69 to build As-negative control |

**Table S6 | Other strains, plasmids and synthesized DNA constructs used in this study**

| Strain, plasmid or construct | Source | Description |
|---|---|---|
| bAG10 | Gift from Paulsson lab | *E. coli* MG1655 (CGSC 6300) strain, *attTN7*::$P_{rpsL}$ – *syfp2* |
| NEB 10-beta | New England Biolabs | Host strain for arsenite biosensors, a derivative of DH10B |
| Addgene plasmid #62730 | Gift from Niels Geijsen | Expresses Cre in bacterial cells, $P_{T7}$ – *cre* |
| Addgene plasmid #134405 | Gift from Mary Dunlop | Fluorescent reporter for Cre activity, $P_{W4}$ – *loxP* – terminator – *loxP* – *mrfp1* |
| T – $P_{W4}$ – *lox511* – TT – *loxP* | Twist Bioscience, contains TT sequence from Chen *et al.* (9) | Synthesized DNA fragment sequence: AAGCTTTGATCAATGACGAACAATAAGGCCTCCCTAACGGGGGGCCTTTTTTATTGATAACAAAACACAGATAAAAAAAATCCTTAGCTTTCGCTAAGGATGATTTCTATCCAATTATTGAAGGCCGCTAACGCGGCCTTTTTTTGTTTCTGGTCTCCCGATTATCAAAAAGAGTATTGCATTAAAGTCTAACCTATAGGAATCTTACAGCCATCGAGAGGGACACGGCGAAATAACTTCGTATAGTATACATTATACGAAGTTATCAAACGCATGAGAAAGCCCCCGGAAGATCACCTTCCGGGGGCTTTTTTATTGCGCACCTCGGTACCAAATTCCAGAAAAGAGACGCTTTCGAGCGTCTTTTTTCGTTTTGGTCCACATAACTTCGTATAGCATACATTATACGAAGTTATTTTAAGAAGGAGAATTC |

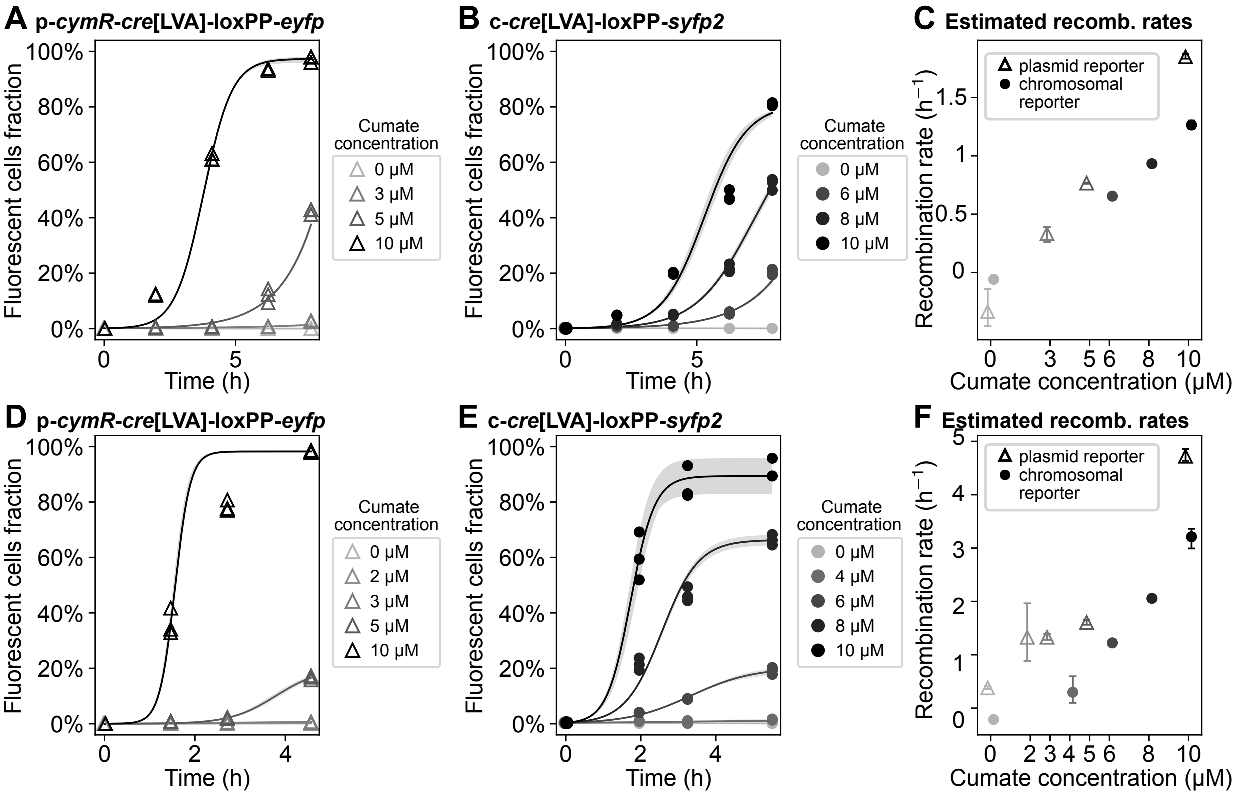


**Fig. S1** Fluorescent cell fractions and recombination rate estimates from flow-cytometry data for additional biological replicates (cf. Fig. 3). Panels A-C and D-F show biological replicates 2 and 3, respectively. Biological replicates 1 (Fig. 3) and 2 (panels A-C) were acquired several months after replicate 3 (panels D-F), using a different 100 mM cumate stock. This deviation may account for the observed differences in recombination rates between biological replicates 1-2 and replicate 3. Each point represents a technical replicate. Time point 0 h is the same for all treatments and is the fraction of fluorescent cells in the starting culture after exiting the lag phase and before adding the inducer. Solid curves show the mean of the best-fit model predictions (Materials and Methods) across technical replicates; shaded bands indicate the min-max envelope of the replicate-specific best-fit curves. Panels C and F show estimated recombination rates obtained via the logistic growth curve fit described in Materials and Methods. Error bars show the minimum and maximum estimated rates across replicates.

In Fig. S1D-F, the following technical replicates were excluded from the analysis due to technical errors, as detailed below:

For the chromosomal reporter c-*cre*[LVA]-loxPP-*syfp2*:

- At time point 5.5 h, a pipetting error occurred during flow-cytometry sampling of one technical replicate induced with 10 μM of cuminic acid.

In addition to the technical replicates shown in Fig. S1D-E, additional technical replicates were initialized and pooled for colony PCR measurements at different timepoints (Fig. S2).

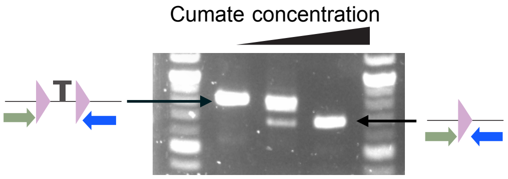


**Fig. S2** Colony PCR of p-*cymR*-*cre*[LVA]-loxPP-*eyfp* exposed for 21 hours to 0, 3 and 10 μM cumate respectively. Gel electrophoresis of PCR products resulting from the amplification of the region enclosing the *loxP* sites using nucleotides oEG48 and oEG102. The gel was exposed to visible blue light for 1 ms. The unrecombined construct yields an amplicon containing two *loxP* sites, whereas Cre-mediated recombination produces a shorter amplicon containing a single *loxP* site. Gel electrophoresis, together with Sanger sequencing of the PCR products, was used to determine whether samples contained unrecombined DNA, recombined DNA, or a mixture of both.

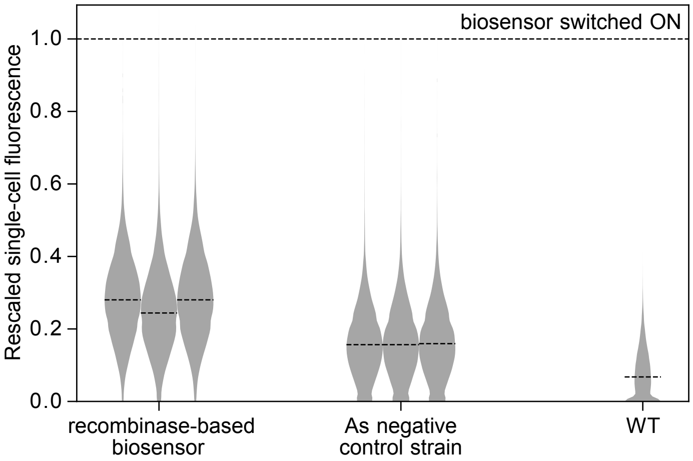


**Fig. S3** Rescaled single-cell fluorescence distributions of overnights of the recombinase based biosensor p-ArsRBS2-*cre*[LVA]-loxPP-*syfp2* (3 biological replicates), As-negative control (3 biological replicates), and the plasmid-free wild-type strain (1 biological replicate), grown in MGP in the absence of arsenic. The background fluorescence of the As biosensor is higher than the negative control, consistent with leaky expression from the $P_{arsR}$ promoter in the absence of inducer. The dashed line in each violin plot shows the median fluorescence of the population. Values are rescaled by the median fluorescence of recombined cells harboring the biosensor plasmid (line at y=1), as a proxy for a positive control.

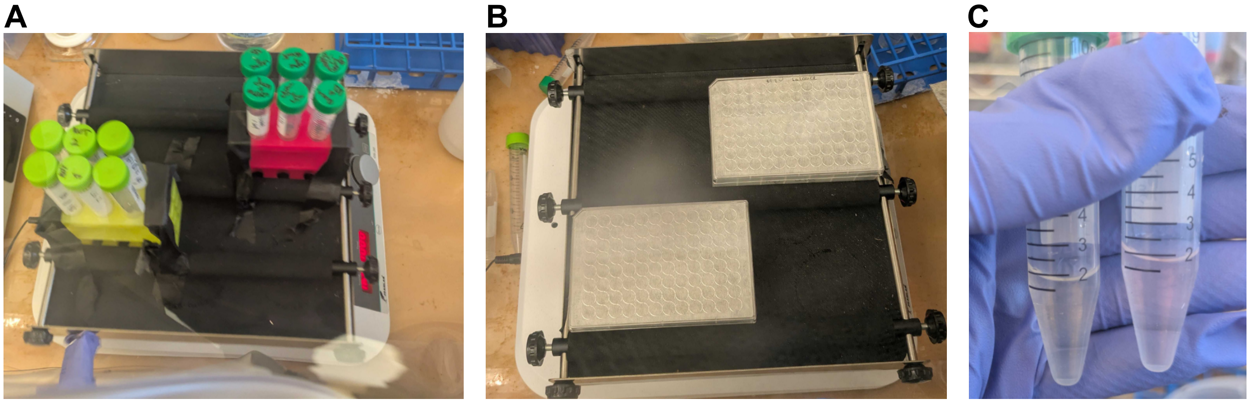


**Fig. S4** Experimental setup for the anoxic arsenite biosensor experiments. Liquid cultures grown on a shaker at room temperature in the anaerobic chamber, before (A) and after (B) arsenite treatment. (C) We validated anoxic conditions of the cell growth medium, by adding 0.001% resazurin. Under aerobic conditions, resazurin undergoes oxidation, resulting in a pink coloration (right tube), while in anoxic environments, it remains in its reduced, colorless form (left tube).

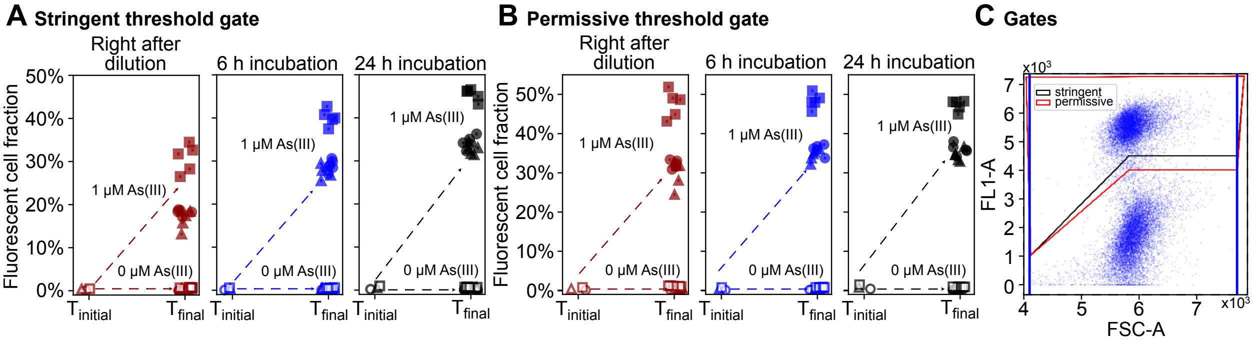


**Fig. S5** Fraction of fluorescent cells before and after anoxic incubation of the arsenite biosensor, with and without arsenite exposure. Fractions were calculated from cytometry data collected at different time points after resuspension of the recombinase-based biosensor in aerobic growth conditions in arsenite-free medium. Different symbols indicate different biological replicates and each point is a technical replicate, with error bars representing the binomial error. Panel A shows fluorescent cell fractions estimated with stringent FACS gating. Panel B shows the same fractions, estimated with less stringent FACS gating that includes cells of moderate fluorescence. Panel C shows the stringent and permissive gates.

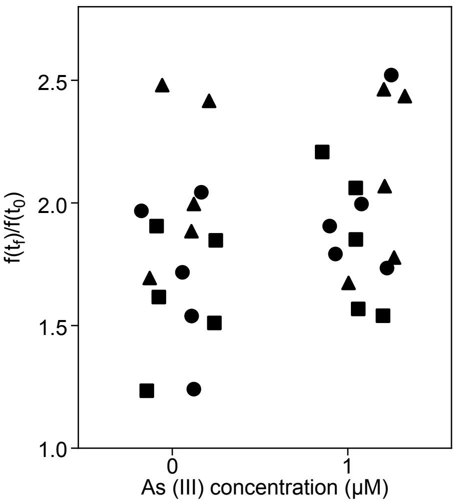


**Fig. S6** Quantification of the fraction of dead cells $f(t_f)$ following anoxic growth, divided by the fraction of dead cells $f(t_0)$ at the start of the experiment. Dead cells were identified via propidium iodide (PI) staining as described in Materials and Methods. PI was added to cells diluted in PBS before measurement at the flow cytometer. This measurement took place right after cells were removed from the anaerobic chamber. Although we observed an increase in the fraction of dead cells during the anoxic growth phase, there was no significant difference in viability between cells grown in the presence or absence of arsenite (*p*-value 0.41). Different symbols represent different biological replicates.

### Additional notes

#### Attempts to clone a cumate-sensitive construct with homologous *loxP* sites and without Cre degradation tag repeatedly led to uninduced Cre-mediated recombination.

Attempting to obtain a construct with homologous *loxP* sites and no Cre degradation tag, we attempted the removal of ssrA-LVA from p-*cymR*-*cre*[LVA]-loxPP-*eyfp* and p-*cre*[LVA]-loxPP-*eyfp*, by PCR-amplifying these plasmids with suitable primers and reassembling them via FastCloning. Alternatively, we used as template a previously assembled circuit on a plasmid already lacking the ssrA-LVA tag, but with heterologous *lox* sites (p-*cymR*-*cre*[LVA]-lox511P-*eyfp*), which we tried to revert to the homologous *loxP* configuration. The modifications of p-*cymR*-*cre*[LVA]-loxPP-*eyfp* and p-*cymR*-*cre*[LVA]-lox511P-*eyfp* attempted in parallel would have returned the same final construct, if successful. The large majority of the transformants across five attempted transformations were fluorescent after overnight incubation on LB agar plates or LB agar supplemented with salicylic acid, which represses $P_{cymRC}$ transcription. Sanger sequencing confirmed that fluorescent transformants had only one *loxP* site and revealed that non-fluorescent ones were false transformants with incorrect plasmid sequences.

### Data analysis

Cumate-sensitive reporter response characterization

Fig. 2 data:

In Fig. 2C, which shows plate reader data for p-*cre*[LVA]-loxPP-*eyfp*, the curves reported for the treatments with 2 μM and 4 μM are the medians computed between 2 biological replicates (corresponding to 6 technical replicates), instead of 3. Samples for the third biological replicate were excluded, since the inducer stock was accidentally diluted beyond the target concentration.

Arsenite-sensitive reporter response characterization

Fig. 4 data:

The following samples were excluded from the analysis (Fig. 4):

- Biological replicate 2 (orange points): Cross-contamination between one technical replicate of pEG68 and one technical replicate of p-ArsRBS2-*syfp2* without arsenite in one well. These two technical replicates were excluded from the analysis, and thus only 4 technical replicates remain. One technical replicate of p-ArsRBS2-*syfp2* (Fig. 4B) treated with 15 nM arsenite did not grow and was thus excluded from the analysis.
- Biological replicate 3 (green points): One technical replicate of p-ArsRBS2-*syfp2* (Fig. 4B) treated with 45 nM arsenite did not grow and was thus excluded from the analysis.

Fig. 5 data:

- The 254 nM arsenic treatment for biological replicate 2 was excluded from the analysis due to a technical issue during sample handling.

Fig. 7 data:

For the cumate-sensitive plasmid reporter, in the last transfer one technical replicate per each condition and biological replicate were not propagated due to a pipetting error. Therefore, those samples were removed from the last measurement.

For the arsenite-sensitive reporter, some samples were excluded from the analysis:

- Biological replicate 1: one technical replicate with 0 nM As was not propagated in the last transfer.
- Biological replicates 2 and 3: for each biological replicate, one technical replicate with 0 nM As was accidentally not propagated in the last transfer; another technical replicate treated with 500 nM As showed an anomalous scatter plot at the flow cytometer, uncharacteristic of *E. coli*, indicating a possible contamination. Therefore, it was removed from the analysis.

**Threshold gates used in cytometry analysis**

Threshold gates were defined manually through the package's graphical interface. For the fluorescent reporter, a gating region was drawn on the FSC-A vs FL1-A (excitation 488 nm, emission 533/30 nm) plot, with the lower boundary set based on the control strain. Dead cells stained with propidium iodide (PI) were gated on the FSC-A vs FL3-A (excitation 488 nm, emission ≥670 nm long pass) plot.
